# Dissecting signaling regulators driving AXL-mediated bypass resistance and associated phenotypes by phosphosite perturbations

**DOI:** 10.1101/2023.10.20.563266

**Authors:** Marc Creixell, Scott D. Taylor, Jacqueline Gerritsen, Song Yi Bae, Mingxuan Jiang, Teresa Augustin, Michelle Loui, Carmen Boixo, Pau Creixell, Forest M White, Aaron S Meyer

## Abstract

Receptor tyrosine kinase (RTK)-targeted therapies are often effective but invariably limited by drug resistance. A major mechanism of acquired resistance involves “bypass” switching to alternative pathways driven by non-targeted RTKs that restore proliferation. One such RTK is AXL whose overexpression, frequently observed in bypass resistant tumors, drives both cell survival and associated malignant phenotypes such as epithelial-to-mesenchymal (EMT) transition and migration. However, the signaling molecules and pathways eliciting these responses have remained elusive. To explore these coordinated effects, we generated a panel of mutant lung adenocarcinoma PC9 cell lines in which each AXL intracellular tyrosine residue was mutated to phenylalanine. By integrating measurements of phosphorylation signaling and other phenotypic changes associated with resistance through multivariate modeling, we mapped signaling perturbations to specific resistant phenotypes. Our results suggest that AXL signaling can be summarized into two clusters associated with progressive disease and poor clinical outcomes in lung cancer patients. These clusters displayed favorable Abl1 and SFK motifs and their phosphorylation was consistently decreased by dasatinib. High-throughput kinase specificity profiling showed that AXL likely activates the SFK cluster through FAK1 which is known to complex with Src. Moreover, the SFK cluster overlapped with a previously established focal adhesion kinase (FAK1) signature conferring EMT-mediated erlotinib resistance in lung cancer cells. Finally, we show that downstream of this kinase signaling, AXL and YAP form a positive feedback loop that sustains drug tolerant persister cells. Altogether, this work demonstrates an approach for dissecting signaling regulators by which AXL drives erlotinib resistance-associated phenotypic changes.

**One-sentence summary:** A systems biology approach elucidates the signaling pathways driving AXL-mediated erlotinib resistance in lung cancer.

## INTRODUCTION

Lung cancer is the leading cause of cancer mortality, accounting for almost 25% of all cancer deaths in the United States for 2022 (*1*). Comprehensive genomic sequencing and expression profiling of non-small cell lung cancer (NSCLC) patients, the most common form of lung cancer, has helped to identify genetic alterations that drive disease progression and can be therapeutically targeted with improved clinical efficacy and safety compared to conventional chemotherapy. One such therapy is the EGFR tyrosine kinase inhibitor (TKI) erlotinib which is effective in patients with advanced EGFR mutant (EGFRm) NSCLC (*2*). However, despite being initially effective, targeted therapies invariably result in incomplete responses, with tumor relapse upon acquiring drug resistance. A major source of resistance to EGFR targeted therapy arises from secondary mutations in the kinase domain, such as the “gatekeeper” EGFR^T790M^, leading to an increased affinity towards ATP relative to its TKI affinity. Second- and third-generation EGFR TKI therapies, such as afatinib and osimertinib respectively, have been developed to effectively target resistance derived from mutant forms of EGFR (*3*). However, while agents targeting EGFR secondary mutations delay tumor relapse, the efficacy of these therapies is ultimately limited by resistance through still other mechanisms.

Another well-appreciated means of resistance to EGFR inhibition is receptor tyrosine kinase (RTK) “bypass” resistance, wherein alternative pathways are activated so that cells are no longer reliant on the drug-targeted pathway. Our lab and others have shown that, while individual RTKs are able to activate a common set of downstream pathways, they do so to varying extents and thus have varied capacity to confer bypass resistance (*4–6*). Two well-studied RTK bypass resistance mechanisms are Her3 signaling providing resistance to Her2-targeted therapy in breast carcinoma and Met signaling driving resistance to EGFR-targeted therapies in lung carcinoma (*5*, *7–9*). Bypass resistance-conferring RTKs may contribute to intrinsic or acquired resistance, can become activated by ligand-mediated autocrine or paracrine induction, amplification, or mutations, and can sometimes be targeted by combination therapy^10^.

AXL, a member of the Tyro3, AXL, and MerTK (TAM) RTK family, is frequently upregulated in tumors resistant to chemotherapy, targeted therapies, and immunotherapy, across cancer types, including EGFR-driven NSCLC (*10–20*). AXL expression is additionally associated with additional phenotypic changes in resistant cells, including epithelial-to-mesenchymal transition (EMT) and cell migration, indicative of increased metastatic capacity (*21–27*). Furthermore, AXL has been shown to sustain the viability of osimertinib resistant cells in lung cancer *in vitro* and *in vivo* EGFRm lung cancer models (*28*, *29*). A landmark study describing AXL-driven erlotinib resistance showed that tumor xenografts with acquired resistance harbored altered expression of several EMT marker genes (*14*). AXL additionally drives resistance to EGFR inhibition of cycling drug-tolerant persister (DTP) cells by protecting them against treatment-induced DNA damage through the activation of low-fidelity DNA polymerases, MYC activation, and a pyridine/pyrimidine metabolism imbalance (*29*). The triple combination treatment of osimertinib, cetuximab, and an anti-AXL antibody led to cures in mice whose tumors had already acquired resistance to osimertinib (*29*). Based on these observations, ongoing phase I/II clinical trials are testing the clinical benefit of AXL and EGFR inhibitor combinations in EGFRm NSCLC (NCT02729298).

Although drug resistance is commonly quantified as a measure or proxy of cell number, tumor relapse is a multifaceted challenge driven by the development of malignant phenotypes that coordinately promote tumor growth and metastasis (*30*). AXL has been involved in a myriad of biological processes that direct cancer progression, including EMT and metastasis (*21*, *22*, *24*, *31*), inhibition of apoptosis (*32*) and induction of cell proliferation (*11*, *28*, *33*), DNA damage repair (*29*, *34*, *35*), endocytosis (*13*, *36*), and tumor immunosuppression (*10*, *20*, *37*, *38*). Consequently, it is unclear exactly which pathways AXL activates to promote resistance, and therefore where, when, and how AXL-mediated resistance develops. Increasing our mechanistic understanding of how AXL regulates these pathways will inform the design of more precise targeted therapy combinations that simultaneously inhibit AXL as well as effector downstream pathways driving resistance. Moreover, a better characterization of AXL signaling might help to better understand the shortcomings of anti-AXL therapies currently undergoing clinical trials. However, identifying these pathways is hindered by RTK crosstalk and signaling pleiotropy: Each RTK regulates a set of downstream pathways that can be also regulated by other RTKs alone or in combination. Therefore, we require a methodology that specifically addresses RTK redundancy and signaling pleiotropy to mechanistically characterize AXL-mediated bypass resistance.

Mutational studies perturbing phosphorylation sites can break RTK pleiotropy into pathway-specific effects. Upon interaction with its cognate ligand, an RTK dimer auto-phosphorylates tyrosine (Y) residues within its kinase domain and C-terminal tail, creating binding sites for interacting adapters and kinases with phospho-binding domains (PBD) such as SH2 domains (*39*). The RTKs then phosphorylate these signaling proteins to initiate downstream signaling cascades (*6*). Since phenylalanine (F) is a non-phosphorylatable mimetic of Y, Y-to-F mutational studies provide a tool for dissecting the signaling role of individual phosphosites. For instance, AXL drives resistance to cetuximab and radiation through interaction of Abl1 with AXL Y821 in head and neck cancer; accordingly, editing Y821 to F abolishes this resistance (*40*). Other AXL Y-to-F mutational experiments have shown that Y821 serves as a docking site for PLCγ, p85, GRB2, as well as Src and Lck (*41*). As individual AXL Y-to-F mutations have different signaling effects, we hypothesized that these mutations would also distinctly mediate AXL’s ability to confer erlotinib resistance, providing an opportunity to establish specific AXL signaling and phenotypic associations during bypass resistance.

Here, we apply an integrated experimental and computational strategy to systematically elucidate the downstream signaling proteins and pathways driving cell survival and associated phenotypes in response to AXL activation. To do so, we generated a cell line panel wherein each cell line carries a single Y-to-F mutation in AXL. We measured phosphorylation signaling alongside viability, apoptosis, migration, and an erlotinib-induced clustering effect in AXL-activated cells treated with erlotinib. Through statistical modeling to analyze the paired AXL-driven phosphoproteomic and phenotypic measurements, our results indicate that AXL activates Src-family kinases (SFK), Abl1, and focal adhesion kinase 1 (FAK1) which serve as key upstream components of the YAP pathway to promote cancer cell growth. Accordingly, we observed that AXL activation drives increased YAP nuclear translocation, whereas YAP inhibition led to decreased AXL expression and kinase activity. The fact that the AXL downstream signaling signature was strongly correlated with AXL expression in EGFRm tumors, as well as poor survival and progressive disease in LUAD patients, supports the clinical relevance of our findings.

## RESULTS

### AXL’s capacity for erlotinib resistance varies across AXL Y-to-F mutants

To isolate the AXL pathways driving specific phenotypic effects, we generated a panel of AXL Y-to-F mutant cell lines and collected phosphoproteomic and phenotypic measurements of each during AXL-driven bypass signaling. We selected these AXL tyrosines based on selection-based FACS assays wherein we transiently introduced an AXL-IRES-GFP plasmid in PC9 AXL KO cells treated with erlotinib (E) and quantified E-based selection (**Fig. S1A/B**). We knocked-out (KO) AXL in PC9 cells using CRISPR/Cas9 and then expressed either AXL wild-type (WT) (KI), AXL kinase-dead (K567R, KD), or one Y-to-F AXL mutant cassette among Y634F, Y643F, Y698F, Y726F, Y750F, or Y821F using lentiviral transduction (**Fig. 1A**). We confirmed AXL total abundance, cell-surface abundance, and activation in each cell line (**Fig. S1C/D**). We observed that around 25% of the transduced PC9 cells express AXL, yet the total AXL expressed and measured by western blot is at least equal to PC9 parental. This suggests that each transduced AXL-expressing cell possesses on average 4-fold more AXL compared with parental cells. While this is a caveat of our methodology, the higher AXL expression of the PC9 Y-to-F mutant cells is still within the range of the reported AXL expression in other EGFRm lung cancer cell lines such as H2279 and H4006 (*27*). AXL activation is challenging to faithfully manipulate given its dependence on phosphatidylserine (PS) present in apoptotic bodies and the spatially heterogeneous presentation of its ligand Gas6 (*42*). Therefore, we used the AXL-activating antibody AF154 (A) as a reproducible means of driving receptor activation. We did not detect AXL phosphorylation in AXL KO and AXL Kinase Dead (KD) cells whereas KI and the Y-to-F mutants showed different levels of kinase activity among each other. Notably, KI and the Y-to-F mutants display lower activation levels than PC9 parental cells which indicate that despite the increased AXL expression in PC9-transduced cells, the total amount of activated receptor on the cell surface is still higher in the parental (**Fig. S1D**). As expected, we observed complete inhibition of p-EGFR upon E treatment (**Fig. S1E)**, and A-induced AXL activity was confirmed by phosphorylation increase of Akt S473-p, a known indirect downstream effector of AXL signaling (*11*, *43–45*) (**Fig. S1F**).

**Figure 1.**
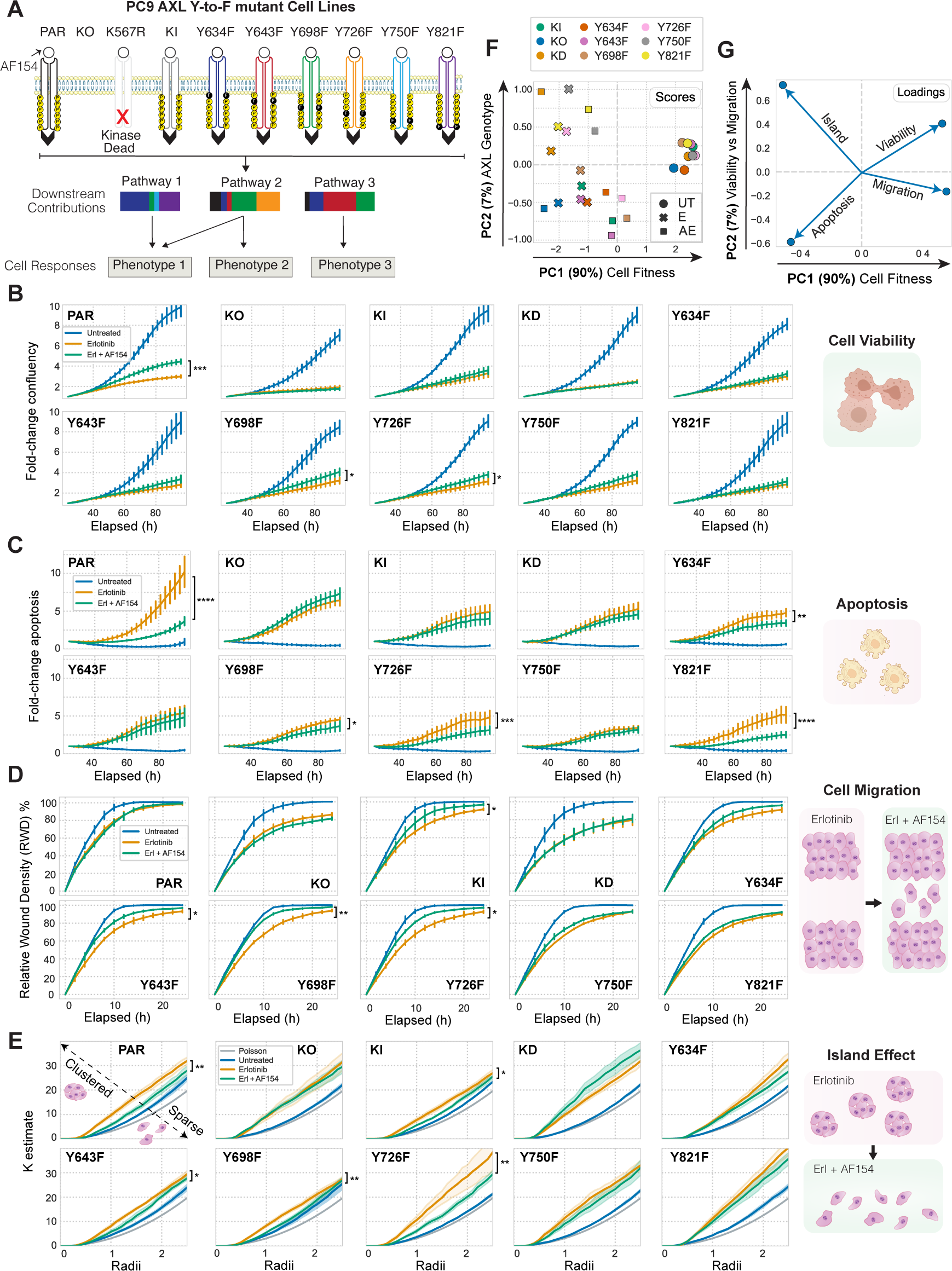
Oncogenic phenotypes vary across PC9 AXL Y-to-F mutants. (A) Schematic of the AXL Y-to-F mutant cell lines each causing distinct signaling and phenotypic consequences upon treatment with erlotinib for 4h and an AXL-activating antibody AF154 for 10 minutes. (B–C) Cell proliferation and cell death quantified for 96 hours using live cell imaging in response to E or EA. (D) Relative wound density (RWD) measured by a scratch wound assay across all PC9 cell lines treated with E or EA. (E) Extent of a E-induced cell island effect upon AXL activation measured by Ripley’s K function. Statistical significance of cell viability was calculated by t-tests in E-versus EA-treated cells across all time points in cell viability, cell death, and migration measurements, and across radii in cell island K estimates. *p-value < 0.05, **p-value < 0.001, ***p-value < 0.0001, ****p-value < 0.00001. Error bars are defined by the standard error of the mean.

Next, we measured the ability of the different AXL mutant cell lines to proliferate, survive, migrate, and form cell “islands” in the presence of E or the combination of E and A (EA) (**Fig. 1A**). We monitored cell proliferation and cell death using live cell imaging over 96 hours of treatment with either E or EA (**Fig. 1B/C**). We observed a significant increase in cell proliferation and decrease in apoptosis in EA-treated PC9 parental cells compared to the E condition. KI cells followed the same trend with lesser magnitude, likely because each PC9 AXL mutant cell line had a lower amount of the receptor on the cell surface than parental (**Fig. S1C**). As expected, EA did not enhance proliferation or survival versus E in AXL KO and KD. While the mutants Y643F and Y750F effectively behaved like AXL KO or KD, Y698F and Y726F cells promoted proliferation and prevented apoptosis upon AXL activation. On the other hand, when EA-treated, Y821F and Y634F cells failed to increase cell proliferation and yet significantly blocked apoptosis (**Fig. 1B/C**).

We performed a scratch-wound assay in parallel to evaluate the migratory capacity of each cell line after treatment. After making a wound, we used live cell imaging for 24 hours to quantify the ability of each cell line to migrate and re-occupy the space in the presence of E or EA. PC9 parental cells treated with E unexpectedly migrated similarly in the presence or absence of the AXL-activating antibody, whereas the migratory ability of KO and KD was blocked, regardless of treatment. Moreover, Y643F, Y698F, and Y726F became more motile after receptor activation, while the mutations Y634F, Y750F, Y821F did not respond (**Fig. 1D**).

While evaluating the other phenotypes, we observed that E induced a clustering effect wherein cells establish cell-to-cell adhesions and form small “cell islands” (**Fig. 1E**). Thus, we asked whether the different PC9 AXL mutant cell lines varied in demonstration of this phenotype. To quantify the extent of cell clustering, we applied Ripley’s K function, a spatial clustering metric frequently used in astronomy to model whether objects such as stars are found closer to one another than would be expected by chance (*46*, *47*). This algorithm allowed us to test the spatial distribution of PC9 cells against the null hypothesis that the cells are distributed randomly. In agreement with our initial observation, we found that parental PC9 cells were more clustered than expected by chance in response to E, whereas AXL activation partially reverted this clustering. Y-to-F mutant cell lines displayed a trend counter to that observed in the scratch wound assay: Y643F, Y698F and Y726F effectively behaved like PC9 parental and KI, whereas Y634F, Y750F, and Y821F remained clustered upon EA treatment (**Fig. 1E**).

We used principal components analysis (PCA) to explore the relationships among the four phenotypes we measured (**Fig. 1F/G**). Most of the variation, represented by principal component (PC) 1, prominently separated untreated cells from the other two treatment groups, and partly separated EA from E cells. We took this component to represent overall cell fitness, which is increased by moving positively along PC 1 for all the mutants except KO and KD, where A had no effect (**Fig. 1F, scores**). Phenotypes increased by AXL activation are positively associated with PC 1, while those decreased are negatively associated. By contrast, PC 2 separated AXL-induced viability from migration effects; moving positively along PC 2 indicated an increase in viability and a decrease in apoptosis while migration decreased; cells formed islands in the opposite direction (**Fig. 1G, loadings**). The cell lines showed varying effects with AXL activation: Y726F and Y750F shifted positively along PC 1 and negatively along PC 2 with A treatment, reflecting migration-driven effects; Y634F and Y821F moved positively along PC 1 and PC 2, reflecting cell viability/apoptosis-driven effects.

Overall, these results suggest that each Y-to-F mutant affects cell viability, death, migration, and clustering differently. On the one hand, representing extremes of phenotypic behavior, while Y750F cells blocked all phenotypic effects, behaving like KO and KD, Y698F and Y726F cells behaved like PC9 parental and KI, with little impact on phenotypic outcomes. On the other hand, representing the most varied phenotypic outcomes, Y634F and Y821F mutant cells blocked AXL-induced viability, migration, and scattering, while simultaneously enhancing cell survival in the presence of E.

### AXL mutants selectively disrupt downstream pathway effects

Intrigued by these differential phenotypic outcomes and to address whether differences in downstream signaling drive these differences, we applied a mass spectrometry approach, as previously described (*48*), to measure the signaling effect of AXL activation in the PC9 AXL mutant cells during EGFR inhibition. We treated each PC9 mutant cell line with E for 4 hours and then stimulated AXL for 10 minutes before lysis. Hierarchical clustering of the resulting phosphoproteomic data set showed AXL KO and KD clustered together, as expected, whereas the other cell lines did not display obvious clustering (**Fig. 2A**).

**Figure 2.**
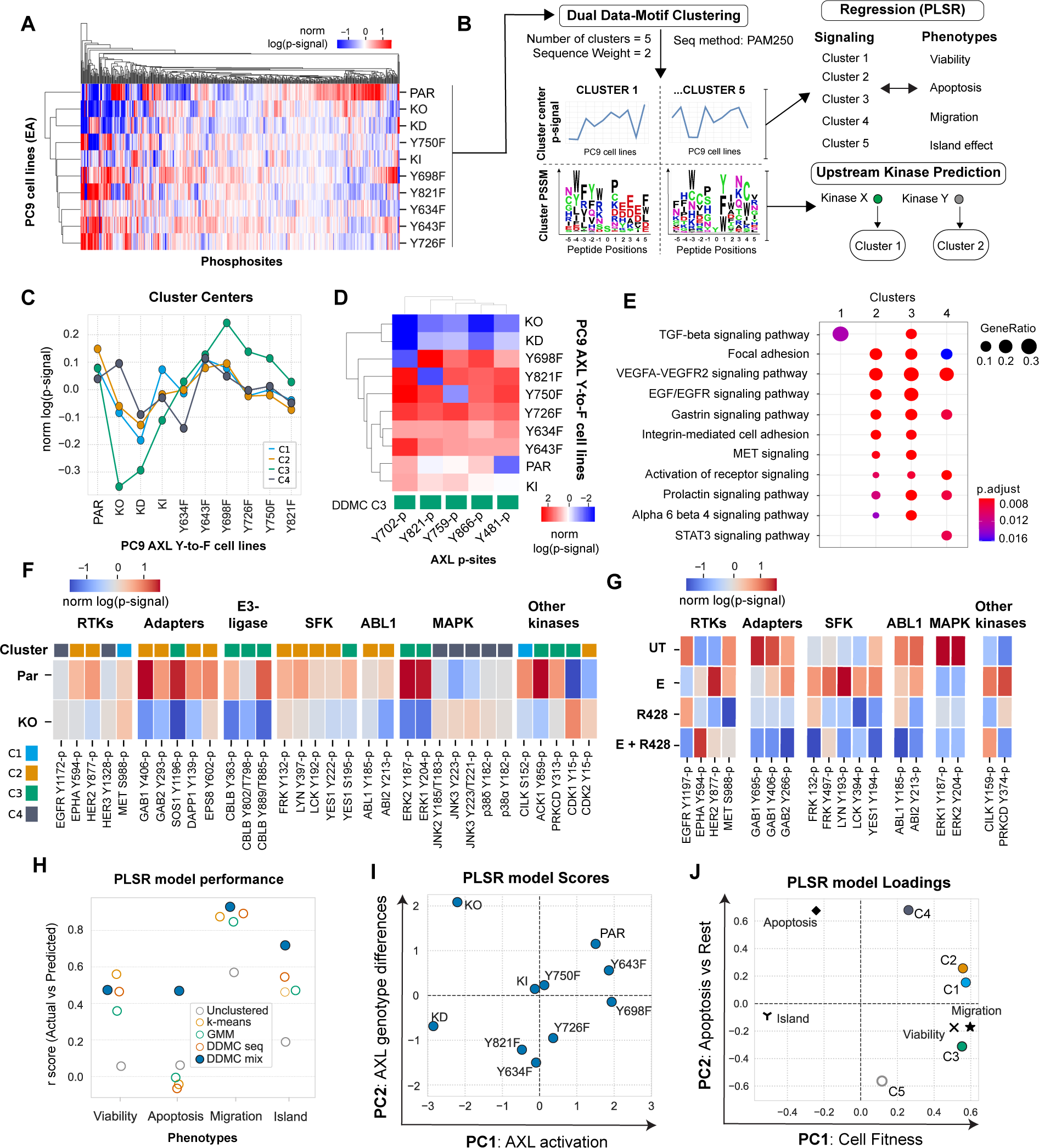
DDMC signaling clusters predict the AXL-mediated phenotypes and identifies CK2, Abl1, and SFK as putative bypass signaling drivers. (A) Global phosphoproteomic measurements from each of the PC9 AXL Y-to-F cell lines. (B) Computational strategy to map the network-level phosphoproteomic changes driving AXL-mediated phenotypic responses. The signaling data was clustered using DDMC to generate 5 clusters of peptides displaying similar phosphorylation behavior and sequence features. The cluster centers were then fit to a PLSR model to predict the phenotypic responses and find associations between clusters and phenotypes. DDMC was used to infer putative upstream kinases regulating clusters. (C) Average relative phosphorylation signal of the DDMC cluster centers. (D) Phosphorylation signal of AXL phosphosites per PC9 cell line and their cluster assignments. (E) Ranked GSEA analysis of DDMC clusters using ClusterProfiler. Gene lists per cluster were ranked based on the log phosphorylation abundance fold change of PC9 parental versus AXL KO cells. (F) Selected phosphosites and their cluster assignments in PC9 parental and AXL KO cells (G) Selected phosphosites in PC9 parental cells treated with E, R428, or both. (H) PLSR model prediction performance using the 5 indicated clustering strategies: no clustering (directly fitting the phosphoproteomic data, 5 cluster centers generated by k-means, clusters from a Gaussian Mixture Model (GMM), DDMC using only the peptide sequence information, or DDMC equally prioritizing the sequence and phosphorylation information. (I-J) PSLR scores (I) and loadings (J).

To identify significant clusters in our data, we applied Dual Data-Motif Clustering (DDMC), an approach that we previously developed to improve the computational modeling of signaling networks by clustering phosphoproteomic data based on both abundance variation and sequence information (*49*). We then used Partial Least Squares Regression (PLSR) to establish associations between these signaling clusters and cellular response and DDMC to infer the upstream kinases regulating each cluster (**Fig. 2B**).

DDMC requires selecting a series of hyperparameters (namely cluster numbers, sequence weights, and PLSR components), which we did by assessing the ability of PLSR to predict the phenotypes using each set of hyperparameters. We additionally inspected the interpretability of the top 5% performing models and selected the one comprised by 5 clusters, a sequence weight of 2, and 2 PLSR components (**Fig. S2A–D**). Several models using both the phosphorylation abundance and sequence information, with a sequence weight of 1, 2, 3, or 5, outperformed all models using either information source exclusively (**Fig. S2C**).

The resulting cluster (C) centers grouped the AXL-responsive behaviors. C1, C2, and especially C3 were markedly decreased in PC9 AXL KO and KD cells, with varied phosphorylation signal across Y-to-F mutants. C3 peptides were of particular interest to us because, in addition to a dramatic phosphorylation decrease in AXL KO and KD, it showed an increased abundance in PC9 AXL Y698F—a mutant that phenotypically behaves like PC9 parental (**Fig. 1**)—compared with the other clusters. Conversely, C4 shows an increase in PC9 KO—but not KD—with respect to parental, and a lower phosphorylation signal in Y634F compared to the rest of clusters (**Fig. 2C**). Cluster 5 had biologically inexplicable signaling trends, and so was disregarded (**Fig. S2E**). We next investigated the cluster membership for all AXL phosphorylation sites (phosphosites) and found all of them within C3, consistent with C3 displaying the most dramatic AXL responsiveness (**Fig. 2C/D**). The lack of correlation between the phosphorylation differences observed in the AXL phosphosites with total or cell surface AXL levels per cell line suggests that the variation in signaling was not due to varying ectopic AXL expression (**Fig. S1C**).

We ran Gene Set Enrichment Analysis (GSEA) on each cluster to explore their functional roles. We found that C1 is strongly enriched in a TGF-beta signaling pathway signature; C2 and C3 share the enrichment of several biological processes associated with the activation of RTK signaling (EGFR / VEGFR) and the regulation of focal adhesions; and C4 is uniquely enriched in a STAT3 signaling pathway signature (**Fig. 2E**). These results highlight known AXL-mediated processes, such as downstream EGFR pathway re-activation and focal adhesion regulation, as well as less established programs such as TGF-beta and STAT3 signaling^27^. To obtain a general view of the composition of the different clusters, we next explored the cluster assignments and phosphorylation of RTKs, receptor adapters, and canonical protein kinases in PC9 parental and AXL KO cells treated with EA (**Fig. 2F**). Among RTKs, we found a decrease in the phosphorylation of Epha2 Y594-p and Her2 Y877-p in AXL KO compared to PC9 parental. On the other hand, we observed an E-induced phosphorylation increase in Met S988-p and Her2 Y877-p in E-treated PC9 parental cells which was abolished in the presence of the AXL inhibitor R428 (**Fig. 2G**). This E-induced phosphorylation is of particular interest because it is consistent with previous studies that have suggested signaling crosstalk between AXL and HER2 to drive resistance to anti-HER2 therapy in breast cancer, as well as between AXL and EPHA2 or MET to confer resistance to EGFR inhibitors (*17*, *28*, *50*, *51*). Of similar interest is the fact that the RTK adapters Gab1/2, Eps8, Sos1, or Dapp1 as well as the E3-ubiquitin ligase Cbl-b are markedly downregulated in AXL KO, and that the phosphorylation of these proteins is dependent on the availability of AXL Y821, as evident from the fact that these phosphosites were not phosphorylated in the AXL Y821F cell line (**Fig. S2F**). Moreover, we observed increased phosphorylation of the Src-Family Kinases (SFK) Frk, Lyn, Lck, Yes1, as well as Abl1 and its substrate and regulator Abi2 in C2 and C3. While Frk, Lyn, and Lck phosphorylation was increased upon E treatment, monotherapy, or concomitant addition of the AXL inhibitor R428 decreased their phosphorylation. R428 also inhibited Abi2 and Abl1, suggesting an AXL-specific activation of SFK and Abl1 (**Fig. 2F/G**). AXL activation led to a strong phosphorylation increase of Y187-p and Y204-p located in the activation segment of Erk1 and Erk2, respectively. AXL-mediated Erk1&2 activation, which are both C3 members, have been previously reported as drivers of erlotinib resistance (*14*). Administration of E (in the absence of the AXL-activating antibody) and R428, as single agents or in combination, inhibited both Erk1&2 phosphosites, which suggests an activation of ERK signaling by AXL. Conversely, JNK2 and JNK3 phosphosites were modestly more abundant in AXL KO than in PC9 parental and clustered in C4 (**Fig. 2F/G**). Given the signaling behavior and cluster composition of C3, we decided to use STRING to map the protein-protein interactions while observing the phosphorylation changes of PC9 parental compared AXL KO cells within C3. We found that in addition to Erk1/2, other kinases are highly regulated by AXL activation within C3 such as Cdk1 (as shown by the lack of phosphorylation of its inhibitory phosphosite Y15), Prkcd, Cilk1, Ack1, and the phosphatase Shp2 (**Fig. S2G**). GSEA indicated that signaling cluster C3 displays increased phosphorylation abundance focal adhesion and cytoskeletal remodeling regulators (**Fig. 2E**). This focal adhesion signature includes SHP2 Y542-p, SIPRA Y496-p, Tjp1 S1196-p, Tjp2 Y249-p, p130cas (*BCAR1*) Y234-p, Actin1 Y215-p, and Afadin (*MLLT4*) Y1230-p, among others (**Fig. 2E**).

In conclusion, the phosphorylation-based signaling landscape varied across AXL phosphosite mutants, demonstrating that single Y-to-F mutations lead to distinct AXL-specific signaling perturbations and cell responses (**Fig. 2 and 3**). DDMC summarized the network-level phosphorylation changes across PC9 Y-to-F mutants during switched AXL activation and found that C1, C2, and especially C3 correlate with AXL activation levels, while peptides in C4 are slightly increased in AXL KO cells compared with parental. These results led us to identify specific signaling-to-phenotype relationships by PLSR.

**Figure 3.**
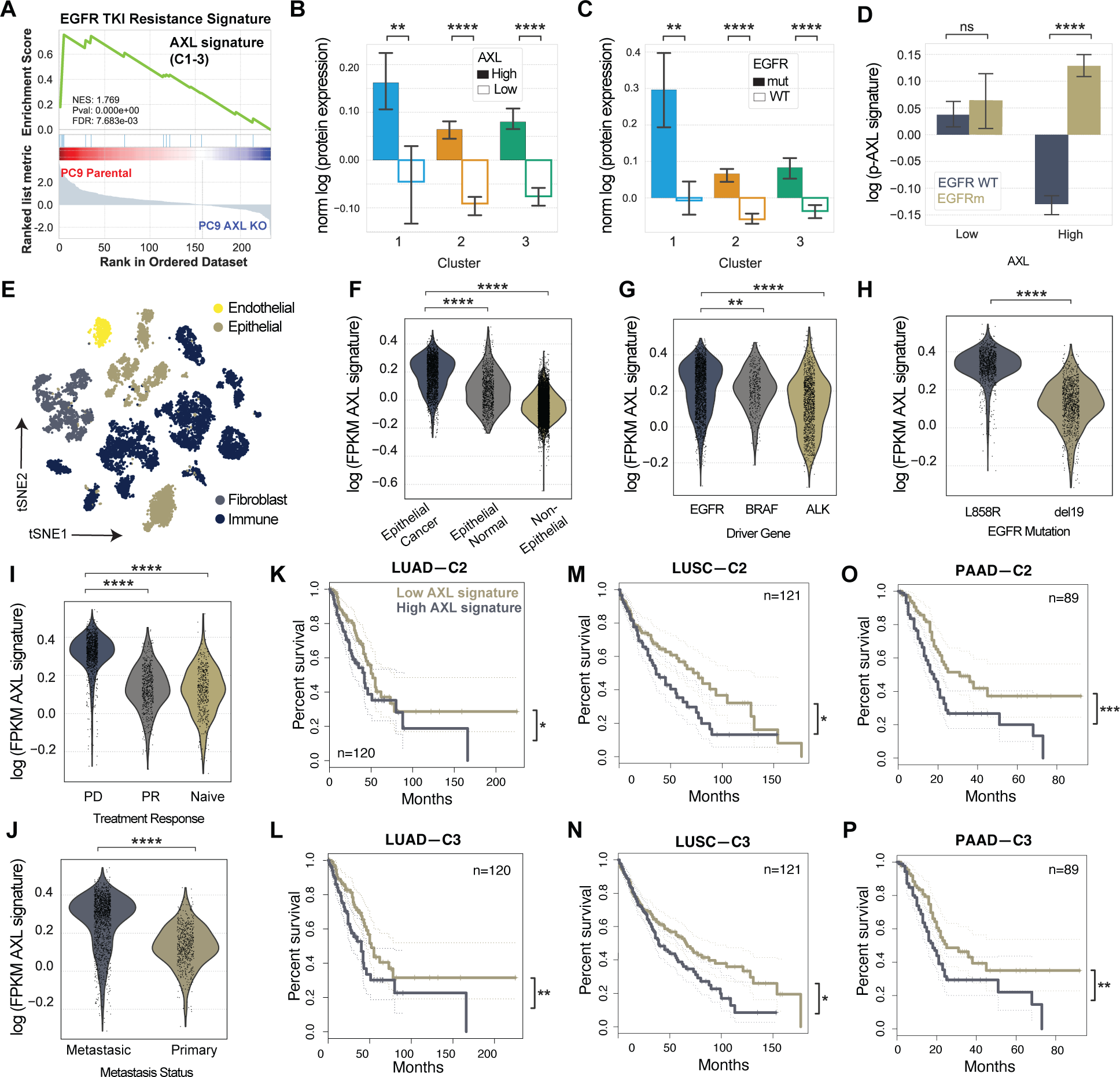
AXL downstream signature based on C2 and C3 is specific to AXL-high EGFRm LUAD tumors and correlates with progressive disease. (A) EGFR TKI resistance signature found by a ranked GSEA analysis using the list of gene names included in C1, C2, and C3 and ranked by their log fold-change phosphorylation between PC9 parental and AXL KO cells. (B–C) Protein expression of C1, C2, and C3 members in (B) AXL-low versus AXL-high tumors or (C) EGFRm versus EGFR WT tumors. (D) Phosphorylation signal of AXL downstream signature by AXL levels and EGFR genotype. (E) tSNE plot of the different cell types of LUAD patient samples defined by Louvain clustering. (F) AXL signature score as defined by the mean gene expression of C1, C2, and C3 per cell in cancer cells, epithelial normal cells, or non-epithelial cells. (G–J) AXL signaling score of cancer cells by (G) driver mutation, (H) EGFR mutation, (I) treatment response or (J) metastatic status. (K–P) Kaplan-Meier curve of (K–L) LUAD, LUSC (M–N), and PAAD (O–P) patients according low or high C2 or C3 gene expression. PD: Progressive disease, PR: Partial Response. PAAD: Pancreatic adenocarcinoma. Error bars in (B-D) show the standard error of the mean. Statistical significance was calculated by Mann-Whitney U rank tests in (B-J) and by logrank tests in (K-P). *p-value < 0.05, **p-value < 0.001, ***p-value < 0.0001, ****p-value < 0.00001, ns means not significant.

### DDMC clusters predict AXL-mediated phenotypic responses and identify C1, C2, and C3 as downstream drivers of erlotinib resistance

To associate the different AXL-mediated signaling clusters and phenotypes, we regressed the DDMC cluster centers against the phenotypic responses using PLSR (**Fig. 2B**). To verify the importance of DDMC-mediated clustering for prediction, we assessed the prediction performance of a PLSR model using different clustering strategies: no clustering (i.e., regressing the phosphoproteomic data set directly), k-means clustering, clustering with a Gaussian Mixture Model (GMM), DDMC using only the sequence information (DDMC seq), or DDMC equally prioritizing the phosphorylation abundance and peptide sequences (DDMC mix). We found that PLSR was only able to accurately predict all four phenotypes when using the cluster centers generated by DDMC mix. By contrast, a PLSR model fit to the MS data directly was not able to predict any of the phenotypes, likely due to the overwhelming number of peptides (498) compared to cell lines (10) (**Fig. 2H**).

The scores and loadings of the resulting PLSR model illustrate the varying capacity of the different PC9 cell lines to promote cell proliferation and migration. PC 1 was indicative of overall cancer cell fitness **(Fig. 2I/J**) like PC 1 of the AXL phenotypes PCA analysis (**Fig. 1F/G**). PC9 parental, as well as the AXL mutants Y698F and Y643F, associated with increased cell fitness positively along PC 1, whereas PC9 AXL KO and KD were heavily negatively weighted along the same axis, thus strongly associating with erlotinib-induced apoptosis and the island effect. PC 2 appears to capture variation that is specific to apoptosis as cells with increased apoptotic phenotypes move positively along this axis. The PLSR model captures an inverse association between apoptosis and the PC9 AXL mutants Y634F and Y821F as they are negatively associated with PC 2, consistent with the exclusive phenotypic effect of Y634F and Y821F in reducing apoptosis (**Fig. 1C and 2I**); their PLSR scores were not weighted along PC 1 and therefore not associated with proliferation, migration, nor the island effect. These results highlight that the phosphoproteomic variation of AXL Y634F and Y821F are consistent with signaling changes that drive their ability to exclusively block cell death, without affecting other phenotypes. We found that only C3 strongly associates with cell migration and proliferation along both PC1 and PC 2, whereas C1, C2, and C3 strongly associate with cell migration and proliferation across PC1 but not PC 2 (**Fig. 2I/J**). Altogether, the results of our paired experimental and computational approach suggest that AXL activates C1, C2, and C3 to promote cell proliferation and migration in the presence of E.

### AXL downstream signaling correlates with poor patient survival and progressive disease in EGFRm LUAD patients

We sought to explore the clinical relevance of the AXL downstream signaling identified by our analysis and its association with resistance to EGFR therapy. To do so, we first selected the phosphosites in C1, C2, and C3 and ranked them by log2 fold-change in PC9 parental versus PC9 AXL KO. This was used to define an “AXL downstream signature” that shows an enrichment of an EGFR TKI resistance signature according to ranked GSEA (**Fig. 3A**).

Next, we interrogated proteomic, phosphoproteomic, and transcriptomic data sets from different clinical studies to investigate the relationship between AXL, its downstream signaling defined by C1, C2, and C3, and patient outcomes. We asked if this AXL downstream signature is enriched in AXL-high LUAD tumors and whether it correlates with poor clinical outcomes. To achieve this, we used data from the National Cancer Institute (NCI)’s Clinical Proteomic Tumor Analysis Consortium (CPTAC) lung adenocarcinoma (LUAD) study which covers the proteogenomic data of over 100 treatment-naïve lung adenocarcinoma tumors (*52*). We stratified the CPTAC LUAD patient samples into AXL-high or AXL-low tumors based on the receptor’s protein expression and looked at the overall and phosphorylation abundance of the members of C1-3. At the protein level, there was a significant enrichment in the abundance of C1, C2, and C3 proteins in AXL-high and EGFRm tumors compared to AXL-low and EGFR WT (**Fig. 3B/C**). Among the 491 phosphosites observed in our AXL mass spectrometry data set (**Fig. 2A**), only 110 were measured in CPTAC. After grouping these peptides into C1, C2, and C3, we did not observe a difference in their phosphorylation abundance when comparing AXL-high versus AXL-low tumors, however, in EGFRm compared with EGFR WT tumors, AXL-high tumors displayed a strong phosphorylation enrichment of the AXL signature, specifically in C2 and C3 but not C1 (**Fig. 3D** and **Fig. S3A/B**). Thus, the identified AXL downstream signaling occurs in clinical specimens and is specific to EGFRm tumors.

Even though phosphorylation and transcriptional changes are often discordant, most of the biological information of tumor biopsies at the single cell resolution is comprised by gene expression. Thus, we asked whether the gene expression of those proteins included in the phosphoproteomic AXL downstream signature is higher in EGFRm tumors displaying progressive disease and metastasis, and thus resistance to EGFR-targeted therapies. Using previously published single cell RNAseq (scRNAseq) data (*53*), we observed that AXL expression was significantly increased in progressive disease and metastatic tumors (**Fig. S3C/D**). The gene expression of our phosphoproteomic AXL downstream signature was significantly upregulated in cancer cells compared with other cell types, in L858R EGFR+ tumors, as well as in progressive disease and metastatic tumors (**Fig. 3E-J**). By plotting each cluster separately, we found that these differences can be accounted for by the contribution of genes present in C2 and C3, but not C1 (**Fig. S3E-S**).

Using TCGA data, we explored whether the gene expression of AXL downstream signaling correlates with poor overall survival of LUAD patients. We observed statistically significant decreases in the percent survival of patients with higher gene expression of C2 and C3 gene expression in LUAD, lung squamous cell carcinoma (LUSC), and pancreatic adenocarcinoma (PAAD) tumors (**Fig. 3J/K**). Enrichment of our AXL signature in PAAD patients with poor clinical outcomes is consistent with the fact that many PAAD tumors are driven by mutant EGFR. Moreover, AXL has indeed also been implicated in resistance to TKI in both LUSC and PAAD (*24*, *54*, *55*).

Above we showed that DDMC identified three clusters of phosphosites regulated by AXL activity that correlate with cell viability and migration in vitro. Here, we asked whether these sites, as well as the transcript and protein expression of these proteins, are associated with poor clinical outcomes in different patient cohorts. At the phosphorylation level, we observed that the AXL downstream signature is specifically enriched in EGFRm and not EGFR WT tumors. At the protein level, the expression of C1, C2, and C3 members was higher in EGFRm and AXL-high tumors. Finally, the transcriptional expression of these proteins showed a remarkable association with poor clinical outcomes.

### Dasatinib selectively targets C2 and C3

Given the strong correlation between the transcript expression of the proteins in C2 and C3 identified in our phosphoproteomic model and poor prognosis in LUAD patients, we decided to further investigate the regulation of these clusters by AXL signaling. A feature of DDMC is the construction of position-specific scoring matrices (PSSMs) for each cluster which represent the frequency of each residue across peptide positions. These computationally derived kinase motifs can then be compared against a compendium of 60 experimentally determined kinase motifs to infer the upstream kinases regulating each cluster (*49*) (**Fig. 2B**).

DDMC inferred that CK2, Abl1, and SFK act upstream of C1, C2, and C3, respectively (**Fig. 4A**). C1, the only pS/T cluster, displays a kinase motif characterized by the strong enrichment of an acidophilic C-terminus, a known hallmark of CK2 specificity (**Fig. S2H**) (*56*, *57*). The GSEA results of C1 provides support to this DDMC upstream kinase prediction as CK2 has been reported to be activated by TGFβ treatment and required for TGFβ-induced EMT (**Fig. 2E**) (*58*). CK2 has also been shown to phosphorylate the C1 members Mcm2, Ldlr, and Sptbn1 (*59*). With C2, the PSSM is consistent with an Abl1 kinase motif; the kinase specificity of Abl1 features a proline at position +3 with respect the phosphorylation site, an isoleucine at -1 and an alanine at +1 (**Fig. S2I**). Both C2 and C3 have sequence motif features associated with SFK: C2 has a strong enrichment of acidic residues at position -3 and to a lesser extent at position -2 (**Fig. S2J**). The SFK Yes1 specifically favors glycine, threonine, and tryptophan at position +1 as well as serine, glycine, and methionine at -2, which are all included in the PSSM of C2. On the other hand, C3 displays a strong enrichment of hydrophobic residues leucine and phenylalanine at position +3, serine and threonine at +2, and basic residues at position +4 and +5, which are all specificity drivers of SFK. C3 was also inferred to be regulated by RTK signaling as its motif is highly preferred by several receptors such as Alk, Met, and INSR. These receptors tend to target substrates that present highly hydrophobic C-terminus in addition to hydrophilic residues at -1, which are characteristics of the motif of C3. The association of C3 with RTKs is illustrated by a STRING network map and is consistent with the GSEA of C3 (**Fig. 2E** and **Fig. S2G**).

**Figure 4.**
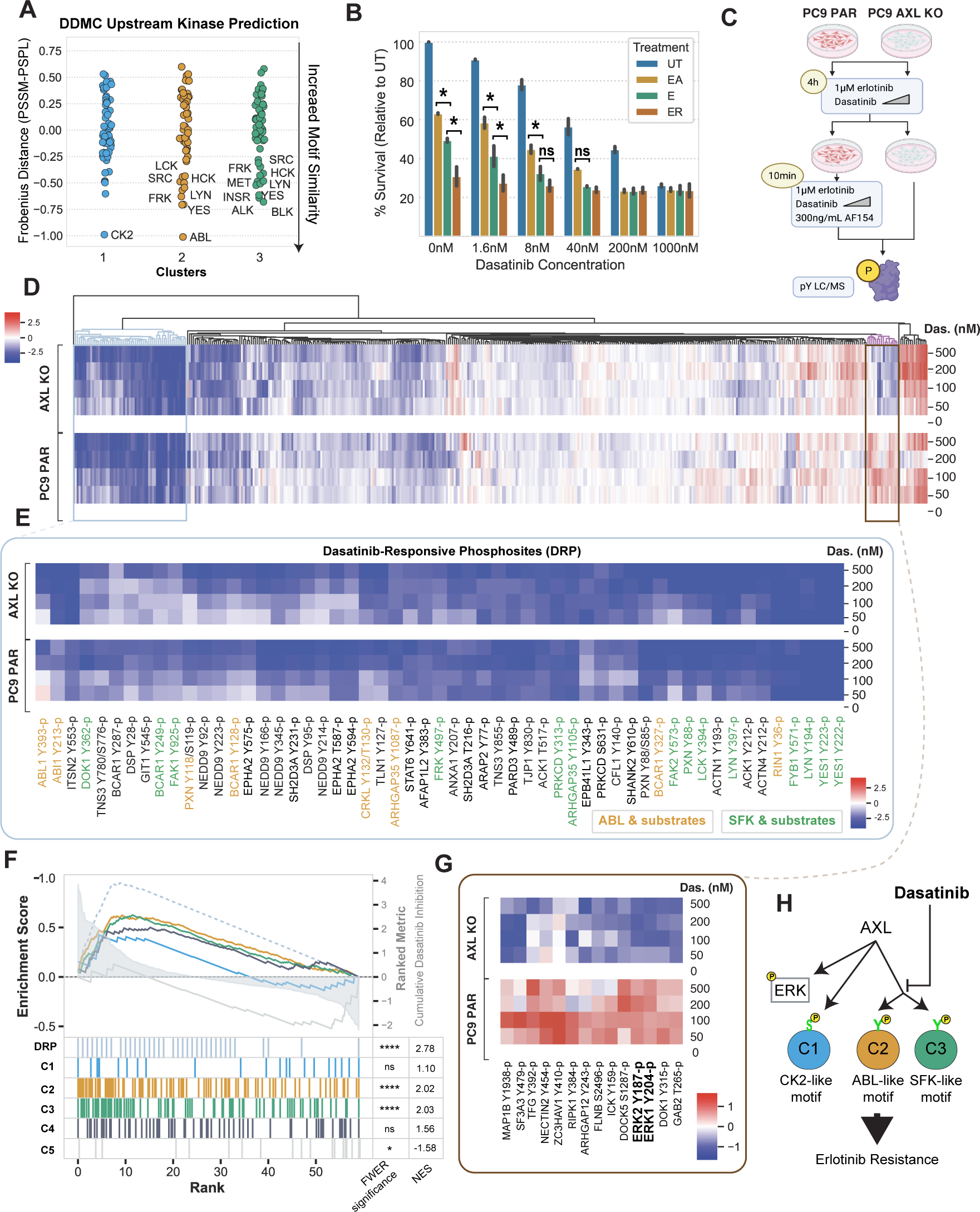
Dasatinib inhibits C2 and C3. (A) DDMC upstream 1475 kinase predictions. (B) Cell confluency of PC9 parental cells exposed to the indicated treatments with increasing concentrations of dasatinib for 72 hours. Data normalized to untreated cells. Statistical significance was calculated by Student’s t-tests. (C) Diagram of the MS experiment. Cells were treated with E and the indicated concentration of dasatinib for 4 hours and subsequently with AF154 for 10 minutes. Cells were then lysed and subjected to mass spectrometry (see Methods). (C) Hierarchical clustering of the entire phosphoproteomics data set of PC9 PAR and AXL KO cells showing the log phosphorylation signal of peptides normalized to the 0 nM dasatinib condition per cell line. (E) Heatmap of dasatinib-responsive phosphosites (DRP). Abl1 and SFK substrates were manually annotated according to PhosphoSitePlus. (F) Ranked GSEA of DRP and DDMC clusters. Peptides were ranked by calculating the cumulative inhibition across increasing dasatinib concentrations. (G) Cluster of phosphosites showing an increased signal in PC9 PAR but decreased phosphorylation in AXL KO cells treated with the indicated concentrations of dasatinib and EA. (I) Cartoon illustrating the effect of dasatinib on AXL downstream signaling. In B and F, *p-value < 0.05, ****p-value < 0.00001, ns means not significant.

To experimentally test these links and whether SFKs and Abl1 may regulate C2 and C3 peptides, we measured cell proliferation and the global phosphoproteomic state of PC9 cells upon treatment with the targeted inhibitor dasatinib. We first asked whether dasatinib was able to block the cell proliferation increase induced by AXL activation. We treated PC9 parental cells with E, EA, or E with the AXL inhibitor R428, all in the context of increasing concentrations of dasatinib. As expected, EA and E+R428-treated cells were significantly more and less proliferative, respectively, than cells treated with E, and we observed a dose-response decrease of cell proliferation with increasing concentrations of dasatinib (**Fig. 4B**).

We performed another pY-based mass spectrometry experiment to ascertain whether there was an enrichment of phosphosites from C2 and C3 in those peptides most prominently depleted by dasatinib treatment. We activated AXL in PC9 parental cells after pretreating with E and increasing concentration of dasatinib for 4 hours. In parallel, we also pretreated AXL KO cells. We then lysed cells and ran mass spectrometry (**Fig. 4C**). Hierarchical clustering identified a cluster of peptides that were highly sensitive to dasatinib (**Fig. 4D/E**). Beyond Abl1- and SFK-related phosphosites, this dasatinib-induced cluster includes most of the AXL-regulated kinases previously described in **Fig. 2** including Ack1, Prkcδ, FAK1/2, and Epha2, as well as proteins associated with focal adhesion regulation and cytoskeletal remodeling such as p130cas, CasL, Git1, Tjp1, Paxillin, and Actin1 (**Fig. 4E**). Highly sensitive peptides to dasatinib had a selective and statistically significant enrichment overlap with C2 and C3 (**Fig. 4F**).

Finally, the dasatinib treatment data revealed an additional cluster of interest that displays AXL-dependent increased phosphorylation (**Fig. 4G**). Erk1 Y187-p and Erk2 Y204-p remained phosphorylated in PC9 parental cells but were strongly inhibited in AXL KO cells (**Fig. 4G**), suggesting AXL-mediated Erk activation is independent of, but affected by, SFK and Abl1. Together, these results indicate that dasatinib abrogates the AXL-mediated survival advantage mainly by inhibiting the phosphorylation of C2 and C3 (**Fig. 4H**).

### A high-throughput bacterial peptide display screen characterizes AXL kinase specificity and identifies FAK1 as a top substrate

Next, we aimed to ascertain which of the identified phosphosites within the bypass signaling cascade might be directly phosphorylated by AXL, thereby differentiating AXL-proximal from AXL-distal downstream signaling. A strategy to address this question is inferring AXL substrates by inspecting its kinase specificity profile and its likelihood of phosphorylating these different phosphorylation sites in vitro. Since a study comprehensively characterizing AXL’s kinase specificity had not been performed to date, we adapted a previously developed high-throughput specificity screen (*60*) to identify AXL substrates and complement our phosphoproteomics analysis. In peptide display screening, a genetically encoded library of peptides is displayed by an eCPX receptor in *E. Coli* and subsequently phosphorylated by incubation with an isolated kinase of interest. Upon phosphorylation, a pan-pY antibody is used to pull down and enrich for phosphorylated peptides, so that by deep sequencing the original and post-enrichment libraries, a per-peptide enrichment score can be calculated. An anti-Myc antibody binding to the c-Myc tag present in all peptides allows the assessment of, and normalization by, the degree to which each peptide is displayed (**Fig. 5A**). An important feature of peptide display compared to previous methods used to measure substrate specificity is that, by encoding specific peptide libraries, we can not only obtain an optimal phosphorylation motif for a given kinase but also assess phosphorylation preference and likelihood for a defined set of peptides. Thanks to this unique feature, here we can accomplish three different goals, namely (i) defining AXL’s optimal substrate motif, (ii) ascertaining whether AXL preferentially phosphorylates the peptides identified in our study relative to most known human phosphotyrosines, and (iii) identifying the highest-scoring substrates within our phosphoproteomics data set.

**Figure 5.**
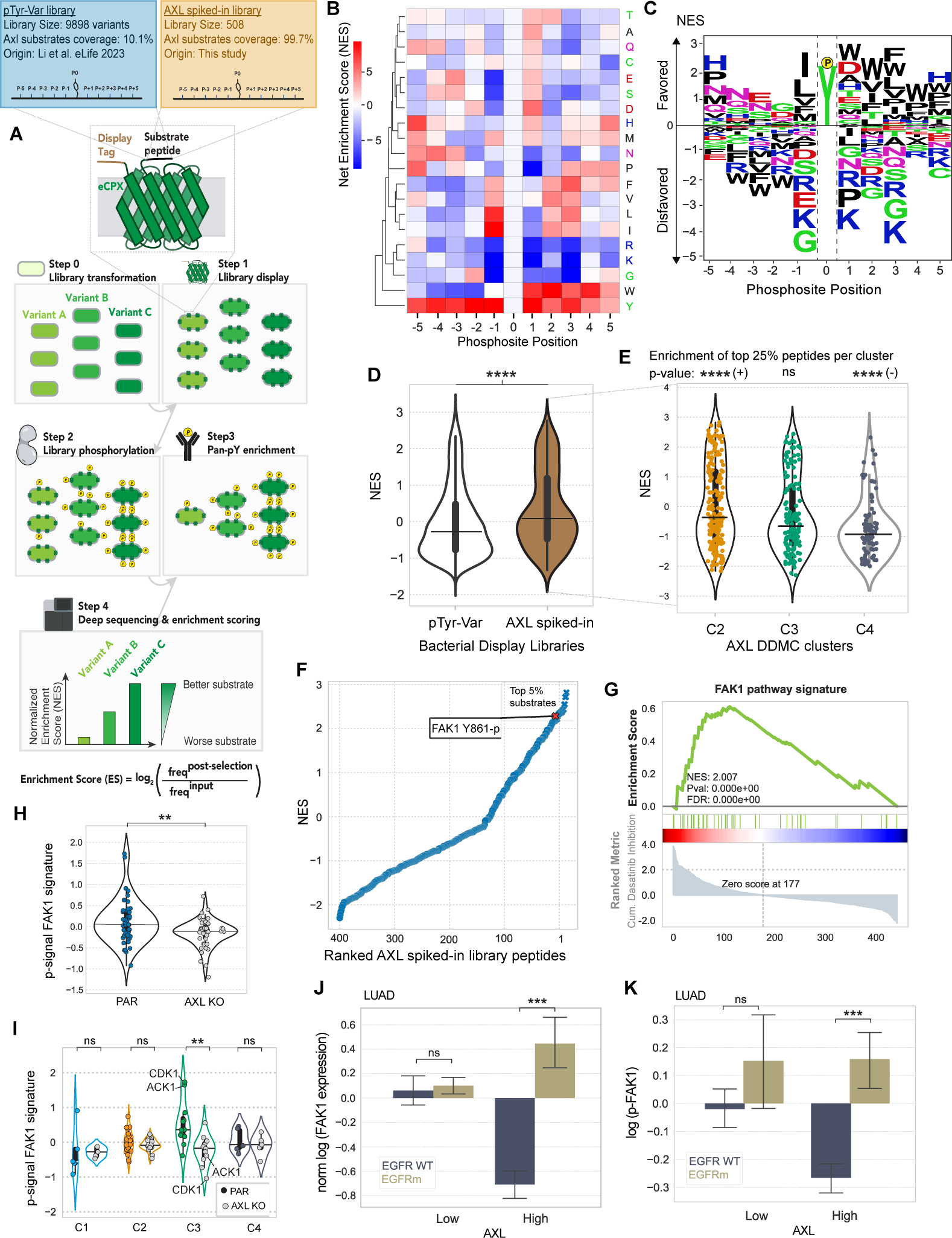
A high-throughput specificity screen shows that AXL directly phosphorylates FAK1 which in turn regulates C3. (A) Schematic describing the screen’s workflow. (B–C) AXL-BTN PSSM illustrated by either a heatmap (B) or a logo plot (C). (D) Violin plot showing the NES distribution split by whether a peptide is present in the AXL phosphoproteomics data set (brown) or not (white). (E) Violin plot showing the NES of all AXL MS peptides grouped by cluster. A hypergeometric test was used to calculate the enrichment of top 25% substrates within each cluster. Signs next to significance markers indicate whether clusters are enriched (+) or depleted (-) with top 25% substrates. (F) Ranked AXL substrates by NES. Refer to **Supplemental Figure 5E** for a selected list of top substrates by cluster. (G) Ranked GSEA of phosphosites of proteins included in the FAK1 pathway signature. Peptides were ranked by calculating the cumulative inhibition across increasing dasatinib concentrations. (H–I) Phosphorylation signal of the FAK1 signature members grouped by either cell line (H) or DDMC cluster (I). Note that the signal of CDK1 Y15-p was multiplied by -1 since it is a known inhibitory site of its kinase activity. (J-K) Pan-FAK1 total protein (I) or phosphorylation (K) in LUAD patient samples stratified by AXL-hi or AXL-low. In D, E, and H-K, statistical significance was calculated using Mann-Whitney U rank tests. *p-value < 0.05, **p-value < 0.001, ***p-value < 0.0001, ****p-value < 0.00001, ns indicates not significant. In J-K, the error bars are defined by the standard error of the mean.

Shah *et al* built a library comprised of 2600 annotated human tyrosine phosphorylation sites— which they, and we, refer to as the “pTyr-Var” library (*60*). Consistent with the fact that the “pTyr-Var” library provides only partial coverage of the human phosphoproteome, we found that only ∼10% of the peptides within our AXL phosphoprotoemics dataset were part of the pTyr-Var library. To study all peptides within our AXL phosphoproteomics dataset, we built a second purpose-made library, hereafter referred to as “AXL spiked-in library”, including the 415 pY peptides from our study that populate C2, C3, and C4 (**Fig. 5A** and **S4A**). While we screened both glutathione S-transferase (GST)- and biotin (BTN)-tagged AXL so that we could rule out significant tag-derived effects, both screens led to largely overlapping peptide specificities, as indicated by a Pearson’s coefficient of 0.96 (**Fig. S4B**), with the most noticeable difference being cysteine enrichment unique to AXL-GST (**Fig. 5B/C** and **S4C/D**). Since we reasoned that this cysteine might be mediated by the relatively larger GST tag, and therefore focused on the results from our AXL-BTN screen.

We computed a PSSM representing AXL’s substrate motif by quantifying the relative enrichment or depletion of a given residue along the peptide sequence positions using phosphorylation abundance of each peptide within the library (**Fig. 5B/C**). AXL shows a strong preference for hydrophobic residues in the C-terminal part of the peptide (positions +1 to +5 relative to the phosphorylation sites). It also favors peptides with hydrophilic residues at +1, namely aspartate or histidine. On the N-terminal side, AXL displays a strong preference for isoleucine or leucine at -1, a common substrate specificity hallmark for many tyrosine kinases. Moreover, AXL preferentially phosphorylates peptides with an asparagine in their N-terminal side, particularly in positions -5 through -3. AXL also disfavors substrates containing the basic residues lysine and arginine, albeit shows a marked preference for histidine at positions -5, +1 and +5. Consistent with the expectation that similar kinases are likely to share similar substrate specificities, AXL’s specificity profile is very similar to the reported substrate specificity of another TAM-family kinase, MerTK (*61*).

By comparing the peptide display screening from the two libraries side-by-side, we observed that the phosphosites included in our phosphoproteomics data set are preferentially phosphorylated by AXL compared with most phosphotyrosines within the proteome (**Fig. 5D**). We then asked how high-scoring peptides in the top 25% within the AXL spiked-in library were represented across the DDMC clusters. We observed an enrichment of the top 25% AXL substrates in C2, no enrichment in C3, and a depletion of top substrates in C4 (**Fig. 5E**).^26,27,62^ While we found no SFK or Abl1 peptides to be directly phosphorylated by AXL, we observed the activating phosphosite Y861-p of FAK1 to be among the top 5% of AXL substrates (**Fig. 5F**). FAK1 is a known regulator of focal adhesions and migration, frequently associated with malignancy and metastasis. A previous study defined that a FAK pathway signature in EMT-mediated erlotinib resistant lung cancer cells was counter-acted by dasatinib (*62*). These mesenchymal resistant cells overexpressed many of the proteins that our model identified to correlate with migration and proliferation when differentially phosphorylated upon AXL activation (**Fig. 2F/G**). Thus, we asked whether our AXL downstream signature overlaps with the FAK signature. Consistent with the study that identified the FAK1 pathway to drive EMT-mediated erlotinib resistance, we found this pathway, like C2&C3, to be significantly depleted by dasatinib (**Fig. 5G**). We saw that AXL-activated PC9 cells display an increased phosphorylation of FAK signaling compared with AXL KO, and that this pathway activation is mainly driven by C3 with Cdk1 and Ack1 being most dramatically activated (**Fig. 5H/I**). Moreover, we observed higher protein expression and phosphorylation of FAK1 in EGFRm AXL-high LUAD tumors (**Fig. 5J/K**), which is consistent with the AXL downstream signaling trends described previously in the same dataset (**Fig. 3B-D** and **S3B**).

Among the top-scoring AXL substrates, our in vitro phosphorylation screen identified several known AXL substrates, such as Ack1, SHP-2, CTNND1, and PEAK1 (*26*, *27*, *63*) as well as less established interactors including the cytoskeletal-remodeling proteins TJP2, BCAR1, TNS1, ANXA2/5, TWF1, ITSN2, the kinase DYRK1A/3, and the E3-ubiquitin ligases CBLB and CBLC (**Fig. S4E**). Six different NEDD9 phosphosites were among the top 10% AXL substrates, consistent with the GSEA enrichment of C2 and C3 and previous studies describing the role of AXL in regulating focal adhesions and migration through this interaction (**Fig. 2F** and **S4E**) (*26*). A ranked GSEA analysis was consistent with most preferred AXL substrates preferentially localizing at the cellular membrane (**Fig. S4F**).

Therefore, this bacterial display screen helped us differentiate AXL-proximal or direct from AXL-distal or indirect bypass signaling machinery. By doing so, we found AXL to highly phosphorylate FAK1 Y861-p in vitro. Moreover, FAK1 signaling pathway largely overlaps with C3 and is effectively inhibited by dasatinib. This suggests that FAK1 might play a critical role in regulating C3 signaling; the cluster most affected by AXL across Y-to-F mutants and that shows the strongest correlation with cell migration and viability in our PLSR model (**Fig. 2C/J**).

### AXL drives upstream YAP regulators which feed back to drive AXL expression and activation

To extend our understanding of the mechanism by which the identified phosphoproteomic signaling changes affect AXL-driven phenotypes, we decided to explore the transcriptional changes occurring during switched AXL signaling. We collected RNAseq data of each PC9 AXL Y-to-F cell line treated with E or EA. PCA analysis of the RNAseq data found that PC1 represents AXL activation since the scores of all cell lines shift positively along PC 1 in EA-treated cells compared with E. We generated a ranked gene list based on the PC 1 scores by ranked GSEA analysis and found a YAP signature enriched in AXL-activated cells (**Fig. 6A** and **S5A**). This is consistent with our DDMC upstream kinase predictions of C2 and C3, as Abl1 and SFK phosphorylation of YAP is a well-known mechanism of YAP nuclear translocation and activation (*64–67*). Moreover, several studies demonstrate an association between YAP activation and the development of drug resistance, including in the context of EGFR-targeted therapy in NSCLC cells (*68–72*). Additionally, a recent study shows that FAK1 signaling is the main driver of YAP activation to enable the emergence of drug tolerant persister cells during EGFR-targeted therapies in patient-derived models and in clinical samples (*73*). Our phosphoproteomics and bacterial display data show that FAK1 and its associated signaling is activated by AXL (**Fig. 5F–K**). Together, these insights led us to hypothesize that AXL is a key upstream component of the YAP pathway—namely SFK, Abl1, and FAK signaling—mediating the emergence of drug-tolerant persister cells.

**Figure 6.**
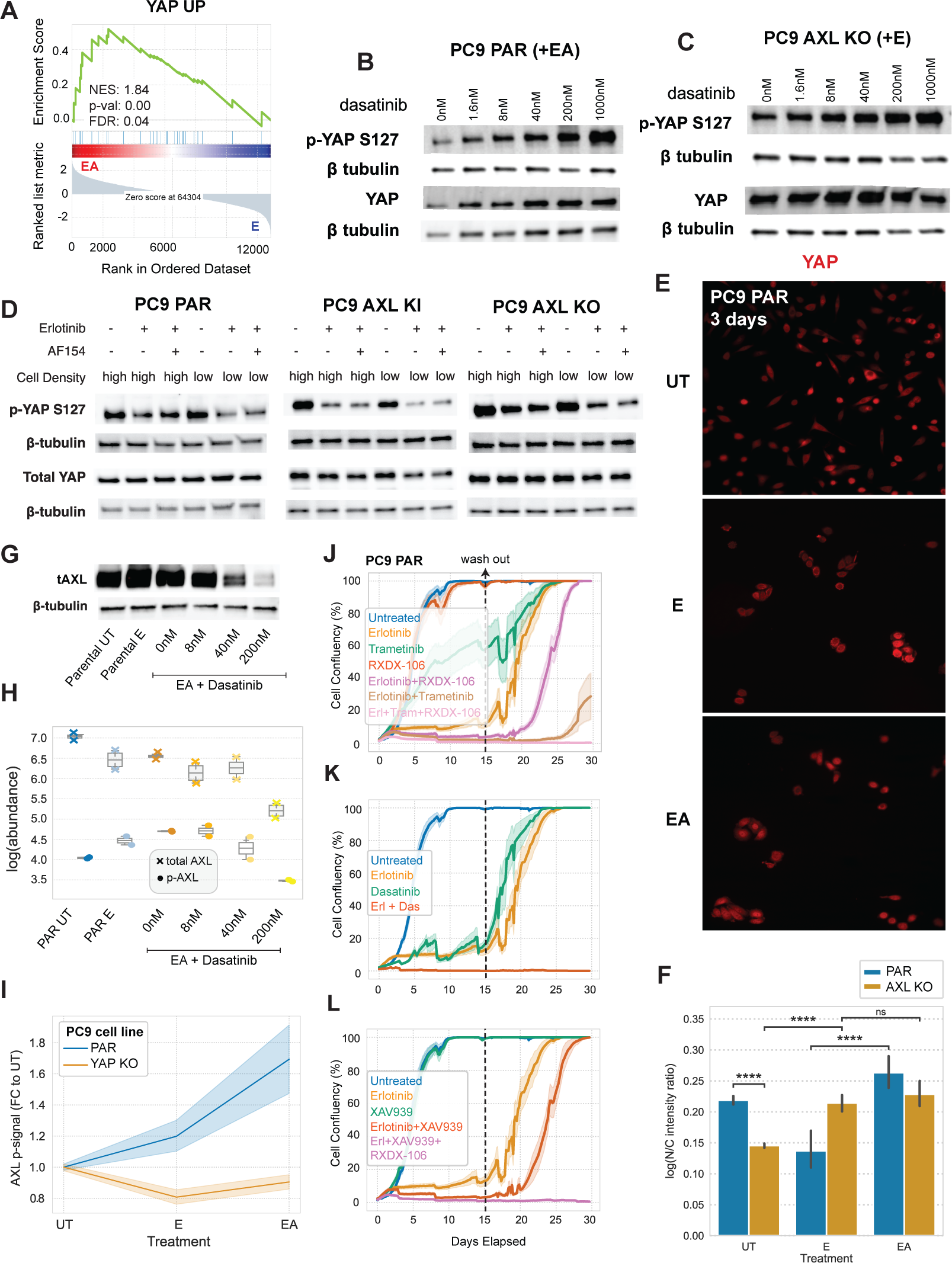
AXL promotes the activation and nuclear translocation of YAP which, in turn, regulates AXL expression and kinase activity. (A) Ranked GSEA analysis of the RNAseq data of the Y-to-F mutant cell lines ranked by the scores of a PCA analysis (see **Fig. S6A**). (B–C) Total and S126-p YAP levels of (B) PC9 PAR and (C) AXL KO cells with increasing concentrations of dasatinib in addition to EA. (D) Total and S126-p YAP levels of PC9 PAR, AXL KI, and PC9 AXL KO cells seeded at high or low cell density and treated with E or EA. (E–F) YAP immunofluorescence staining in PC9 parental cells under the indicated treatments for 3 days (E) and the corresponding quantification including AXL KO measurements (F). Statistical significance was calculated using a Mann-Whitney U rank test. ****p-value < 0.00001, and ns means not significant. (G) Western blot of total AXL in PC9 cells. (H) Luminex of total and phospho-AXL in PC9 cells. (I) Luminex of phospho-AXL in PC9 PAR and PC9 YAP KO cells treated with E and EA. (J–L) Cell viability assay of PC9 PAR cells treated with the indicated inhibitors for 15 days. Treatment conditions were replaced with media and drug-tolerant persister cells were allowed to regrow for 15 days. Treatment or media were refreshed every 3–5 days. All error bars or regions show the standard error of the mean.

To test this hypothesis, we first treated cells with increasing concentrations of dasatinib and blotted for the inhibitory YAP phosphosite S127 which, when phosphorylated, is bound by 14-3-3 proteins to sequester YAP in the cytoplasm and prevent its nuclear translocation (*64*). Indeed, we found that dasatinib induces an increase in the phosphorylation abundance of the YAP S127-p which is further exacerbated in the absence of AXL. AXL KO cells had higher amounts of total YAP protein expression compared with PC9 parental cells (**Fig. 6B/C**).

To further validate the effect of AXL signaling on YAP activation, we measured YAP S127-p in PC9 parental, AXL KI, and AXL KO cells treated them with E or EA for 24h (**Fig. 6D**). We seeded cells at high or low density as YAP is known to respond to changes in cell confluency, with the transcription factor becoming inactive at higher cell densities (*74*). As expected, we found that YAP S127-p most strongly decreases in response to E or EA at low cell density. While AXL KO cells displayed the same trend, the overall phosphorylation signal of S127-p is substantially higher in AXL KO compared to PC9 parental or AXL KI cells, again reflecting AXL-specific YAP activation.

To further assess YAP state, we directly quantified YAP cellular localization in PC9 parental and AXL KO cells using immunofluorescence. We treated cells with E or EA for 3 days to observe YAP localization in cancer cells persisting in the presence of inhibitor. PC9 cells treated with EA display significantly higher nuclear YAP than cells treated only with E (**Fig. 6E/F**). Moreover, while most AXL KO cells have cytoplasmic YAP, parental cells displayed much more nuclear localization of YAP (**Fig. 6F**). Consistent with the western blots of YAP S127-p, we observed AXL-independent, erlotinib-induced YAP nuclear translocation (**Fig. 6E/F** and **Fig. S5B/C**).

While several studies indicate that AXL can be a target of YAP when in complex with the DNA-binding TEAD proteins, thereby proposing that YAP acts upstream of AXL (*68*, *75*, *76*), others have reported that it is in fact AXL kinase activity which induces YAP activation (*72*). A third role emerged when another study showed that AXL and the YAP homologue TAZ form a positive feedback loop to promote the development of lung cancer brain metastases (*31*). Our results additionally suggest that upon AXL activation, SFK, Abl1 and FAK1 kinases engage a network of cytoskeletal-remodeling, EMT-associated, signaling proteins that affect YAP translocation. We therefore explored the effect of indirectly disrupting YAP activity, via dasatinib, on AXL expression and activation and found that both total and phosphorylated AXL in PC9 parental cells decrease with increasing concentrations of dasatinib (**Fig. 6G/H**). Moreover, PC9 YAP KO cells treated with E or EA fail to activate AXL (**Fig. 6I**). Together, these data indicate that AXL and YAP form a positive feedback loop.

We then investigated the influence of AXL signaling in the emergence of drug-tolerant persister cells. We inhibited AXL, Erk1/2, SFK/Abl1/FAK1, or YAP through RXDX-106, trametinib, dasatinib, and XAV-939 treatment, respectively, in PC9 parental, AXL KO, or YAP KO cells for 15 days. XAV-939 is a tankyrase inhibitor that is widely used as an indirect inhibitor of YAP activity (*70*, *77*, *78*). After treatment for 15 days, we replaced the treatment solutions with complete media and assessed the ability of resistant cells to regrow. We used RXDX-106 instead of R428 because RXDX-106 is more AXL-specific; R428 has been shown to trigger AXL-independent cell death (*79*). We found that EGFR inhibition alongside genetic or pharmacological inhibition of AXL modestly delayed the emergence of DTP cells, while combined EGFR and Erk1/2 blockade more strongly inhibited DTP cells (**Fig. 6J** and **Fig. S5D**). Triple inhibition of EGFR, AXL, alongside either YAP or Erk1/2, completely blocked the emergence of resistant cells (**Fig. 6K/L**). Consistent with the notion that dasatinib disrupts both AXL and YAP activity (**Fig. 6B/G/H**), the combination of E and dasatinib prevented cancer cell regrowth (**Fig. 6K**). The same trends were observed in AXL KO cells wherein both combination treatments E and trametinib, E and XAV939, or E and dasatinib blocked DTP cell growth (**Fig. S5D-F**). The fact that PC9 YAP KO cells were unable to develop resistance to E, dasatinib, or any combination treatment emphasizes the importance of YAP pathway activation for survival (**Fig. S5G-I**).

Taken together, this data supports the hypothesis that AXL and YAP form a positive feedback loop wherein AXL activation leads to the phosphorylation of key upstream components of YAP which, in turn, induces its nuclear translocation and effector functions to sustain cancer cell growth and promote drug resistance. Importantly, while the concomitant inhibition of EGFR and AXL, EGFR and YAP, or EGFR and Erk1/2 delayed the emergence of DTP cells, the triple combination treatment of EGFR, AXL and YAP or Erk1/2 completely blocked the ability of PC9 cells to develop drug resistance.

### AXL-high LUAD tumors display strong Yap activation and EMT marker enrichment

To validate the association between AXL signaling and YAP activation in tumors, we examined the proteogenomic data of 110 treatment naïve LUAD patients from the CPTAC (*52*). AXL abundance was modestly increased in EGFRm versus EGFR WT tumors across stages and, within EGFRm patients, AXL increased throughout disease progression (**Fig. 7A**). Stratifying LUAD tumors into the top and bottom 33% AXL abundance revealed a significant increase in the EMT markers *TWIST1*, *VIM*, and *CDH11* in AXL-high compared to AXL-low tumors (**Fig. 7B**). Leveraging the fact that, unlike in proteomics and phosphoproteomics data, all transcripts are measured by RNAseq in all tumor samples, we were able to confidently ascertain whether a Yap signaling signature is associated with AXL protein expression. As expected, we found a strong enrichment of Yap signaling (*80*) in AXL-high tumors by a GSEA analysis wherein tumors were ranked based on their AXL protein expression (**Fig. 7C**). Overall, these results in combination with the scRNAseq analysis of LUAD tumors suggest that some aspects of the AXL-mediated bypass signaling characterized *in vitro* may translate in patients (**Fig. 3** and **7A-C**).

**Figure 7.**
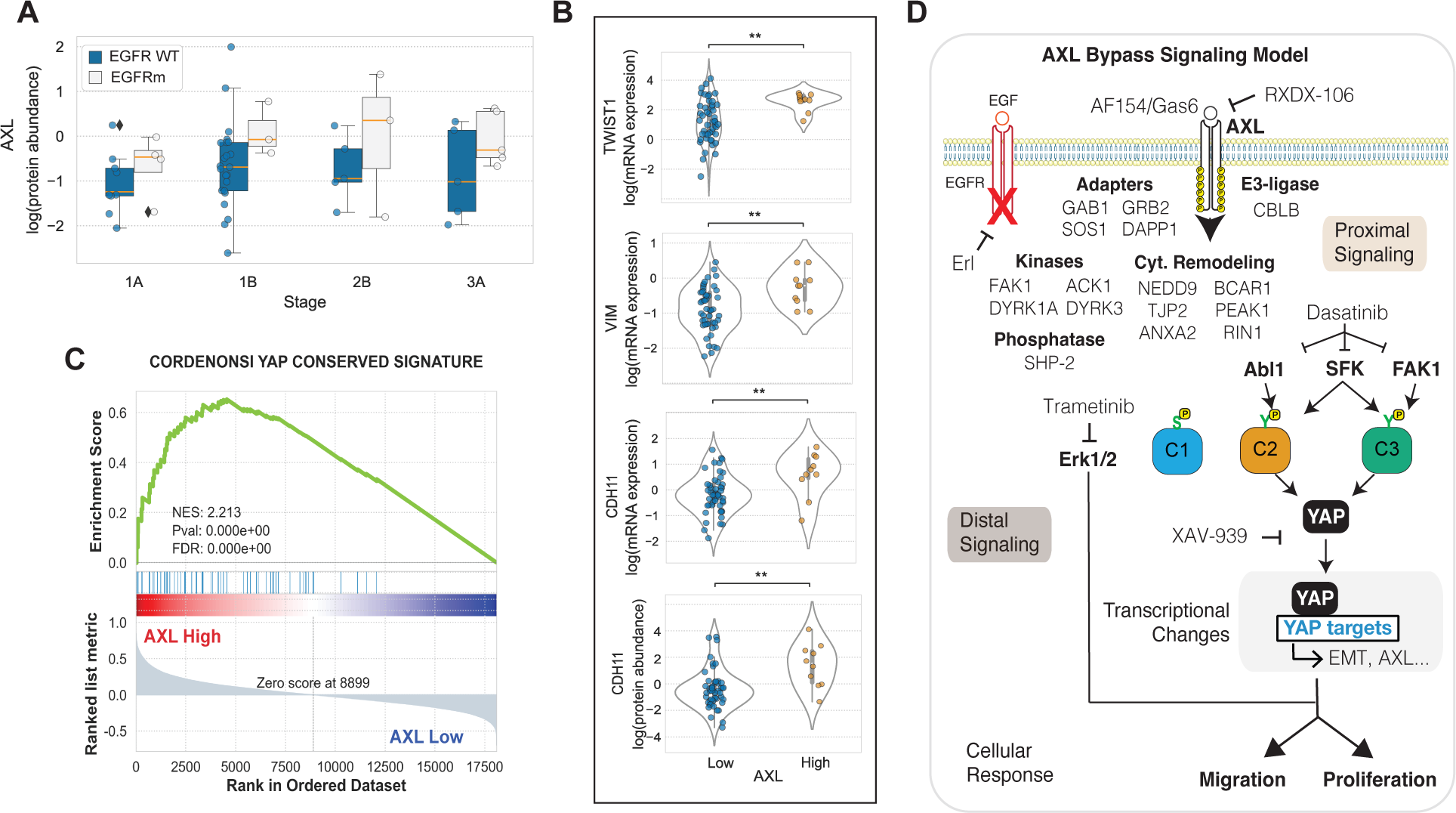
AXL-high LUAD tumors display increased YAP activation and EMT markers. (A) AXL protein levels grouped by EGFR mutational status and tumor stage. (B) Expression of mesenchymal markers by AXL levels. **p-value < 0.001 according to Student’s t-test. (C) Transcriptomic YAP signature in AXL-high vs AXL-low tumors. (D) Illustration of the AXL bypass signaling network identified in this study.

In this study, we propose a kinase downstream signaling model during switched AXL activation in E-treated PC-9 cells (**Fig. 7D**). Our modeling and specificity screening results (**Fig. 2** and **5**) indicate that AXL proximally engages the adapters GAB1, SOS1, GRB2, and DAPP1 through Y821, a signaling cascade of cytoskeletal remodeling proteins such as NEDD9, PEAK1, and TJP2, as well as the kinases FAK1, ACK1, and DYRK1A/3 and the phosphatase SHP-2. Our model identified three clusters that most prominently correlate with malignancy, namely the phospho-serine cluster C1, and C2 and C3 which are formed by phospho-tyrosine sites. Given that we performed tyrosine phosphoproteomics and that, unlike C2 and C3, the transcriptional changes of C1 members did not correlate with poor patient outcomes (**Fig. 3**), we decided to focus on C2 and C3. DDMC inferred that both clusters generally displayed favorable kinase motifs for SFK, whereas C2 additionally included key determinants of Abl1 specificity which suggested to us that these kinases might act as upstream regulators of C2 and C3 (**Fig. 4A** and **Fig. S2H-J**). SFK and Abl1 inhibition via dasatinib treatment effectively blocked AXL-mediated bypass resistance and selectively decreased the phosphorylation of C2 and C3 (**Fig. 4A/F**). Our bacterial specificity results showed that FAK1 Y861-p is a top AXL substrate in vitro and we observed that a FAK1 pathway signature was substantially inhibited by dasatinib treatment (**Fig. 5F/G**), which led us to hypothesize that FAK1 might play an important role in AXL bypass signaling. Indeed, we observed that FAK1 pY signaling was significantly enriched in PC9 parental compared with AXL KO cells and that the differentially phosphorylated peptides pertaining to the FAK1 signature were clustered in C3 (**Fig. 5H/I**). We then asked what transcriptional programs were most affected by AXL bypass signaling and identified a YAP signaling signature (**Fig. 6A**). We then experimentally showed through western blots, Luminex, and immunofluorescence that AXL activation induces YAP nuclear translocation and effector functions which in turn allows AXL activation (**Fig. 6B-I**). Finally, combination treatment of E and the Abl1/SFK inhibitor dasatinib, or the triple combination treatment of E, the AXL inhibitor RXDX-106, and indirect inhibition of YAP via XAV-939 or Erk1/2 inhibition via trametinib blocked the emergence of drug-tolerant persister cells (**Fig. 6J-L**).

## DISCUSSION

The reactivation of oncogenic pathways mediated by RTKs not targeted by therapy, referred as RTK bypass signaling, is a well-known resistance mechanism (*5*, *7–9*). Met activation has been shown to mediate bypass resistance to EGFR-targeted therapies in lung cancer (*7*) and, in fact, the identification of this resistance mechanism led to the recent FDA-approval of Amivantamab, a bispecific antibody directed against EGFR and MET for patients with advanced or metastatic EGFRm NSCLC (NCT04599712) (*81*). However, we still do not have a clear grasp on the signaling landscape that bypass RTKs activate to mediate drug resistance. Identifying key signaling networks driving bypass resistance may lead to (i) an increased mechanistic understanding of RTK inhibitors reaching the clinic and (ii) the identification of new drug targets. Several challenges intrinsic to the study of RTK downstream signaling, such as RTK crosstalk or signaling pathway interdependency, hinder the identification of the specific RTK phosphoproteomic and phenotypic changes driving bypass resistance (*4*, *5*). To overcome this limitation, we devised a combined computational and experimental approach centered around the use of a panel of PC9 AXL Y-to-F mutant cell lines. Using these cell lines, we measured their phosphoproteomic and phenotypic changes during EGFR inhibition and AXL activation and applied multivariate modeling to identify the most prominent AXL-driven signaling pathways that promote cell fitness. This methodology naturally associates molecular and phenotypic variation observed across cell lines to identify AXL- and phenotype-specific signaling.

We identified three phosphosite clusters (C1, C2, and C3) that define an “AXL downstream signature” correlating with cell viability and migration in vitro (**Fig. 3G/H**). C2 and C3, but not C1, correlated with poor treatment response, metastasis, and overall survival in LUAD patients alongside AXL expression. The gene and protein expression of the AXL downstream signature were significantly enriched in AXL-high EGFRm LUAD tumors (**Fig. 3B/C**) whereas the phosphorylation signal was significantly higher in EGFRm LUAD compared with WT tumors (**Fig. 3D** and **S3B**). It remains to be shown, however, whether the identified bypass signaling machinery is conserved across several RTKs or is AXL-specific. A re-implementation of this methodology with a wider panel of RTKs would help elucidate this question and similarly separate the various pathways downstream of each receptor. We observed some amount of AXL-independent, E-induced YAP activation, which suggests that other RTKs might also activate oncogenic YAP signaling. This is consistent with the fact that combined EGFR and AXL inhibition only achieves a delay in the emergence of resistance and that additionally blocking AXL downstream effectors such as YAP, the YAP upstream kinases SFK/Abl1/FAK1, or Erk1/2 completely kills DTP cells (**Fig. 6J–L**). Therefore, our data suggests that concomitantly targeting bypass downstream signaling might provide superior therapeutic efficacy to inhibiting only RTKs.

The Hippo pathway effector YAP is known to drive resistance to EGFR-targeted therapies in NSCLC (*65*, *66*, *69*, *82*). Although YAP has been shown to upregulate AXL gene expression, other studies suggest that it is AXL which acts upstream of the transcription factor (*68*, *72*, *75*). DDMC indicated that C2 and C3 display kinase motifs favored by Abl1 and SFK, respectively, and we found that dasatinib selectively inhibited both clusters, alongside YAP activation, which suggests that these clusters work upstream of YAP (**Fig. 4A and 6B**). Moreover, transcriptomic profiling during switched AXL activation revealed a strong enrichment of YAP targets in AXL-activated cells (**Fig. 6A**). A recent study showed that a previously established FAK1 signaling signature activates YAP to mediate osimertinib resistance, whereas FAK1 inhibition through VS-4718 reduced YAP nuclear translocation and DTP survival (*73*). EA-treated PC9 cells show increased phosphorylation of C3 phosphosites of proteins pertaining to this FAK1 signature compared with EA-treated AXL KO cells which suggests that DTPs, at least in part, rely on AXL for FAK1-mediated YAP activation (**Fig. 5H/I** and **6K/L**).

Whereas previous reports indicate that SFKs and Abl1 can be phosphorylated by AXL (*25*, *31*, *40*), our bacterial display specificity screen shows that neither SFK nor Abl1 motifs are preferentially phosphorylated by AXL in vitro, suggesting that either such phosphorylation is particularly favored in cells or that this link is indirect. On the other hand, FAK1 which is a known regulator of SFK signaling, is among the top 5% of putative in vitro AXL substrates (**Fig. 5F**). FAK1 is known to form a dual kinase complex with Src, and therefore AXL might indirectly activate SFK signaling by promoting FAK1 activity (*62*, *83*). Importantly, genetic ablation of YAP or inhibition of the YAP upstream kinases SFK and Abl1 by dasatinib decreased AXL protein expression and its kinase activity (**Fig. 6G–I**) which indicates that the AXL-SFK/Abl1/FAK-YAP axis forms a positive feedback loop to promote malignancy.

Resistant cells to single-agent osimertinib can emerge through a variety of mechanisms, including Erk1/2 reactivation or YAP activation, whereas YAP activation becomes the dominant resistance mechanism of cells overcoming combined EGFR and MEK inhibition (*70*). Here, we show that AXL can mediate both Erk1/2 reactivation and YAP activation during bypass signaling (**Fig. 2F** and **6**). Intriguingly, we show that SFK/Abl1 inhibition, and consequently YAP inactivation, through dasatinib treatment in EA-treated PC9 cells promotes Erk1/2 activation, whereas dasatinib treatment leads to loss of Erk1/2 phosphorylation in AXL KO cells (**Fig. 4G**). This suggests that AXL might enable cancer cells to rely in either Erk1/2 or YAP to overcome EGFR inhibition. Consistent with this hypothesis, long-term viability assays with TKIs showed that targeting AXL in addition to Erk1/2 or YAP is required to block the emergence of DTP cells (**Fig. 6J–L** and **S5D–I**).

A caveat of our methodology is that each AXL-transduced PC9 cell line expressed about 25% the amount of AXL on the cell surface and around 4-fold more of total receptor compared with their parental counterpart (**Fig. S1C**). The phosphoproteomic and phenotypic measurements from these cell lines were utilized to generate C1, C2, and C3 which allowed us form hypotheses about the key signaling pathways activated by AXL. We then performed computational explorations using proteogenomic data sets of LUAD patients (**Fig. 3**) as well as experimental validations exclusively on PC9 parental and AXL KO cells (**Fig. 4-6**) to corroborate the signaling changes identified by our model. Therefore, we argue that while varying AXL expression in the different cell lines is a limitation of this methodology, the insights generated from this study must be considered holistically, including the follow-on experimental and computational validations.

Altogether, our results suggest that systematically leveraging Y-to-F mutational studies and combining phosphoproteomic with phenotypic experiments allows dissecting pleiotropic signaling regulators to identify the downstream signaling by which AXL drives multiple phenotypes associated with erlotinib resistance in lung cancer.

## MATERIALS AND METHODS

### Antibody reagents, inhibitors, and cell culture

Erlotinib (LC Laboratories), RXDX-106 (Selleck Chemicals), R428 (Fisher Scientific), XAV-939 (Selleck Chemicals) were used at 1 μM, dasatinib (LC Laboratories) at 200 nM, and CX-4945 and trametinib (both from MedChem Express) were used at 4 μM and 30 nM, respectively. The AXL-activating antibody AF154 (R&D Systems) was used at 300 ng/mL. AXL, YAP, YAP S126-p were purchased from CST and rhodamine-conjugated β-tubulin from Bio-Rad and used for Western blotting. ELISA-based signaling measurements were performed according to the manufacturer’s instructions (Bio-Rad). The Luminex kits EGFR Y1068-p and p-AKT is S473-p were obtained from Bio-Rad. The capture AXL antibody was generated by conjugating a primary AXL antibody (R&D systems MAB154) onto magnetic beads (Bio-Rad) as previously reported (*42*). For total AXL detection, a biotinylated AXL antibody (R&D systems BAF154) was used and for p-AXL measurements a pan-tyrosine biotinylated antibody (R&D systems BAM1676) was used. The primary antibodies YAP (Santa Cruz), p-H2AX (CST), and AXL (Abcam), diluted at 1:50, 1:200, and 1:1000, respectively, and the secondary antibodies Alexa Fluor 488 goat anti-rabbit IgG (Invitrogen) and PE goat anti-mouse IgG (Invitrogen), diluted at 1:500 and 1:50, respectively, were used for immunofluorescence as previously described (*42*). PC9 (Sigma Aldrich) cells and all derivatives were grown in RPMI-1640 media supplemented with 10% fetal bovine serum (FBS) and penicillin/streptomycin. HEK293T cells were grown in Dublecco’s Modified Eagle Medium (DMEM) supplemented with 10% FBS and 1% GlutaMAX (Thermo Fisher Scientific). PC9 YAP KO cells were obtained from Passi Jänne’s laboratory at Harvard Medical School. The oxidative stress assay was performed using CellROX^TM^ Deep Red Reagent following the manufacturer’s instructions.

### Generation of PC9 AXL Y-to-F mutant cell lines

The PC9 AXL KO cell line was generated by transfecting cells with a CRISPR/Cas9 and GFP vector containing a gRNA targeting the AXL kinase domain. The gRNA sequence, cell culturing, and sorting methods have been previously described (*84*). Plasmids containing the AXL phosphosite mutations were generated from an AXL-IRES-Puro vector (Addgene #65627) using site directed mutagenesis. Each mutant was then inserted into a lentiviral vector with a puromycin resistance marker (Addgene #17448).

For viral packaging, HEK 293T cells were seeded at 4.5 × 10^6^ per 10 cm dish. After 24 hours, the lentiviral AXL expression vector, VSV-G envelope vector, and packaging vector (Addgene #12259 and #12260 respectively) were combined in a 10:1:10 mass ratio and diluted in Opti-MEM (Thermo Fisher Scientific). TransIT-LT1 (Mirus Bio) was added dropwise, and the solution was mixed gently by swirling and incubated at room temperature for 20 minutes. The solution was then added dropwise to the cells. After 18 hours, transfection media was replaced by media supplemented with 1% BSA fraction V (Thermo Fisher Scientific). Cells were incubated for 24 hours, after which the virus-containing media was removed and stored at 4℃. The media was replaced, and the cells incubated a further 24 hours to generate a second batch of viral media. The harvested batches were then pooled, filtered through a 0.45 μm PVDF membrane to remove packaging cells, and flash frozen followed by storage at -80°C until use.

PC9 AXL KO cells were seeded at 1.5 × 10^5^ cells per well with antibiotic-free media in a 6-well plate and incubated for 24 hours. The cells were then infected with viral particles in antibiotic-free media supplemented with polybrene (MilliporeSigma). After 18 hours, the media was replaced with fresh antibiotic-free media. Cells were observed for a GFP positive population and then passaged into a 10 cm plate until confluent. The virally transduced cells were then sorted for based on GFP expression using a BD FACSAria cell sorter. The mutant cell populations were subcultured for later experiments.

### Cell Viability and Apoptosis Assays

Cells were seeded in a 96-well plate at a density of 1.0 × 10^3^ cells per well. After 24 hours, treatments were added in media containing 300 nm YOYO-3 (Thermo Fisher Scientific). Cells were cultured and imaged every 3 hours using an IncuCyte S3 (Essen Bioscience) at 20x magnification with 9 images per well. The phase, green, and red channels were manually thresholded and then analyzed by IncuCyte S3 software (Essen Bioscience) to determine cell counts and fraction of area covered.

### Cell Migration Assay

96-well IncuCyte ImageLock plates (Essen Bioscience) were coated with a Collagen I solution (Thermo Fisher Scientific), washed twice, and then seeded with 4 × 10^4^ cells per well. After a 4-hour incubation, cells were wounded using the IncuCyte WoundMaker, washed twice to remove detached cells, and then treated with respective conditions. Images of the center of the wound were taken every 2 hours at a magnification of 10x, one image per well. The phase, green, and red channels were manually thresholded and then analyzed by IncuCyte S3 software (Essen Bioscience) to determine migration measurements.

### Cell Island Effect

Phase contrast images used for the cell island measurements were taken from image sets gathered in the cell viability assay. For endpoint readings, images at the 48-hour post-treatment time point were used. Representative images were chosen across experimental replicates. Images were opened in ImageJ and the center of each cell was manually marked. Dead cells, identified using YOYO-3 based fluorescence, were not marked. The 2D coordinates of all cell centers in an image were then exported for analysis. The amount of clustering present in a particular image was then measured by applying Ripley’s K function to the set of coordinates. The implementation of Ripley’s K function used was taken from the astropy Python package (*47*).

### Preparation of Cell Lysates for Mass Spectrometry

Cell lines were grown to confluence in 10 cm dishes over the course of 72–96 hours, washed, and treated by addition of media containing 1 μM erlotinib. Cells were incubated for 4 hours at 37°C and then additionally treated with media containing 1 μM erlotinib and 300 ng/mL AXL activating antibody for 10 minutes. The cells were then placed immediately on ice, washed with ice-cold phosphate-buffered saline, and lysed with cold 8 M urea containing Phosphatase Inhibitor Cocktail I and Protease Inhibitor Cocktail I (Boston BioProducts). The lysates were centrifuged at 20,000xg and 4℃ to pellet cell debris, and the supernatants removed and stored at -80℃. A bicinchoninic acid (BCA) protein concentration assay (Pierce) was performed according to the manufacturer’s protocol to estimate the protein concentration in each lysate. Cell lysates were reduced with 10 mM DTT for 1 hour at 56℃, alkylated with 55 mM iodoacetamide for 1 hour at RT shielded from light, and diluted 5-fold with 100 mM ammonium acetate, pH 8.9, before trypsin (Promega) was added (20:1 protein:enzyme ratio) for overnight digestion at RT. The resulting solutions were acidified with 1 mL of acetic acid (HOAc) and loaded onto C18 Sep-Pak Plus Cartridges (Waters), rinsed with 10 mL of 0.1% HOAc, and eluted with 10 mL of 40% Acetonitrile (MeCN)/0.1% HOAc. Peptides were divided into 200 aliquots, and sample volume was reduced using a vacuum centrifuge (Thermo) and then lyophilized to dryness for storage at -80℃. TMT labeling for multiplexed analysis was performed according to manufacturer’s protocol. Samples, each containing ∼200 μg peptides, were resuspended in 35 μL HEPES (pH 8.5), vortexed, and spun down at 13,400 rpm for 1 minute. 400 μg of a given channel of TMT10plex (Thermo) in anhydrous MeCN, was added per sample. Samples were shaken at 400 rpm for 1 hour, after which the labeling reaction was quenched using 5% hydroxylamine (50%, Thermo). After another 15 minutes on the shaker, all samples were combined using the same pipette tip to reduce sample loss, and sample aliquots were washed twice with 40 μL 25% MeCN/0.1% HOAc, which was added to the collection tube to improve yield. Sample volume was reduced using a vacuum centrifuge and then lyophilized to dryness for storage at -80℃.

### Phosphopeptide Enrichment

Immunoprecipitation (IP) and IMAC were used sequentially to enrich samples for phosphotyrosine containing peptides. TMT-labeled samples were incubated in IP buffer consisting of 1% Nonidet P-40 with protein G agarose beads conjugated to 24 μg of 4G10 V312 IgG and 6 μg of PT-66 (P3300, Sigma) overnight at 4℃. Peptides were eluted with 25 μL of 0.2% trifluoroacetic acid for 10 minutes at room temperature; this elution was performed twice to improve yield. Eluted peptides were subjected to phosphopeptide enrichment using immobilized metal affinity chromatography (IMAC)-based Fe-NTA spin column to reduce non-specific, non-phosphorylated peptide background. High-Select Fe-NTA enrichment kit (Pierce) was used according to manufacturer’s instructions with the following modifications. Eluted peptides from IP were incubated with Fe-NTA beads containing 25 μL binding washing buffer for 30 minutes. Peptides were eluted twice with 20 mL of elution buffer into a 1.7 mL microcentrifuge tube. Eluates were concentrated in a speed-vac until ∼1 μL of sample remained, and then resuspended in 10 μL of 5% acetonitrile in 0.1% formic acid. Samples were loaded directly onto an in-house constructed fused silica capillary column (50 μm inner diameter (ID) × 10 cm) packed with 5 μm C18 beads (YMC gel, ODS-AQ, AQ12S05) and with an integrated electrospray ionization tip (∼2 μm tip ID).

### LC-MS/MS Analysis

LC-MS/MS of pTyr peptides was carried out on an Agilent 1260 LC coupled to a Q Exactive HF-X mass spectrometer (Thermo Fisher Scientific). Peptides were separated using a 140-minute gradient with 70% acetonitrile in 0.2 mol/L acetic acid at flow rate of 0.2 mL/minute with approximate split flow of 20 nL/minute. The mass spectrometer was operated in data-dependent acquisition with the following settings for MS1 scans: m/z range: 350 to 2,000; resolution: 60,000; AGC target: 3 × 10^6^; maximum injection time (maxIT): 50 ms. The top 15 abundant ions were isolated and fragmented by higher energy collision dissociation with following settings: resolution: 60,000; AGC target: 1×10^5^; maxIT: 350 ms; isolation width: 0.4 m/z; collisional energy (CE): 33%; dynamic exclusion: 20 seconds. Crude peptide analysis was performed on a Q Exactive Plus mass spectrometer to correct for small variation in peptide loadings for each of the TMT channels. Approximately 30 ng of the supernatant from pTyr IP was loaded onto an in-house packed precolumn (100 μm ID × 10 cm) packed with 10 mm C18 beads (YMC gel, ODS-A, AA12S11) and analyzed with a 70-minute LC gradient. MS1 scans were performed at following settings: m/z range: 350 to 2,000; resolution: 70,000; AGC target: 3×10^6^; maxIT: 50 ms. The top 10 abundant ions were isolated and fragmented with CE of 33% at a resolution of 35,000.

### Peptide Identification and Quantification

Mass spectra were processed with Proteome Discoverer version 2.5 (Thermo Fisher Scientific) and searched against the human SwissProt database using Mascot version 2.4 (MatrixScience, RRID:SCR_014322). MS/MS spectra were searched with mass tolerance of 10 ppm for precursor ions and 20 mmu for fragment ions. Cysteine carbamidomethylation, TMT-labeled lysine, and TMT-labeled peptide N-termini were set as fixed modifications. Oxidation of methionine and phosphorylation of serine, threonine, and tyrosine were searched as dynamic modifications. TMT reporter quantification was extracted, and isotope corrected in Proteome Discoverer. Peptide spectrum matches (PSM) were filtered according to following parameters: rank=1, mascot ion score>15, isolation interference<40%, average TMT signal>1,000. Peptides with missing values across any channel were filtered out.

### Preprocessing of Phosphoproteomic Data

We performed three phosphoproteomic biological replicates of the PC9 AXL Y-to-F mutants treated with EA. The three data sets were concatenated, mean-centered across cell lines, and log2-transformed. To discard phosphosites whose measurements were not reproducible among replicates, all recurrent phosphosites in two or three biological replicates were identified. For those appearing in two biological replicates, any peptides showing a Pearson correlation coefficient smaller than 0.55 were filtered out, whereas for those appearing in all three biological replicates, peptides with standard deviations of 0.5 or more were discarded. To discard any unchanging peptides across cell lines, we filtered out phosphosites out containing less than a 0.5-fold change between the maximum and the minimum across every cell line. Overlapping peptides were then averaged across replicates. The resulting preprocessed phosphoproteomic data set was then fit using DDMC (*49*).

### Prediction of AXL-mediated Phenotypes using PLSR

To predict the AXL-mediated phenotypes, a 4-component PLSR model (scikit-learn) was built using the DDMC cluster centers. To assess the predictive performance of PLSR, leave-one-out cross-validation was applied; the model was trained using the paired cluster centers and phenotypic measurements in all cell lines except one. The cluster centers of that remaining cell line were used to predict its phenotypic measurements and a mean squared error between the predicted and actual value per phenotype was computed. This process was iterated across all cell lines to obtain a final predictive correlation per phenotype. To benchmark the ability of PLSR to predict the phenotypes using the DDMC clusters, we additionally fit PLSR using either the unclustered phosphoproteomic data set directly, k-means clustering, a Gaussian Mixture Model (GMM), DDMC using only the peptide sequence information, or the selected DDMC model combining the phosphorylation and sequence information (referred as DDMC mix). All cluster methods were used with 5 clusters and the number of PLSR components used was optimized for each case.

### Hyperparameter Selection

The preprocessed phosphoproteomic data set was fit to DDMC using 5 clusters, a sequence weight of 2, and 2 PLSR components. This hyperparameter combination was determined via an exhaustive hyerparameter search (**Figure S3A-D**). Using scikit-learn’s GridSearch, we assessed the ability of the pipeline model comprised of DDMC and PLSR to predict the AXL-mediated phenotypes using different hyperparameter combinations; namely 2 to 15 clusters, a sequence weight of 0, 1, 2, 3, 5, 10, 50, 100, or 500, and 1 to 4 PLSR components. Note that a model with a sequence weight of 0 uses only the phosphorylation abundance for peptide clustering, a weight of 500 mainly uses the sequence information, and intermediate values use a combination of both information sources.

### RNAseq Sample Preparation and Sequencing

To generate the RNAseq data, 300,000 cells of each PC9 AXL Y-to-F mutant cell line were seeded in 100 mm dishes. The next day, each was treated with E or EA and, after 24 hours of treatment, cells were lysed with RIPA and RNA was extracted using the RNeasy Mini Kit (Qiagen). RNA sequencing and read alignment was performed by Novogene. Any genes with less than 10 TPM were filtered out. Ranked or standard GSEA was implemented in Python using the package gseapy (*85*).

### Clinical Data and Analysis

Bulk RNAseq, proteomic, and phosphoproteomic data of LUAD patients was obtained from the CTPAC LUAD study (*52*). scRNAseq of LUAD patients including cell type and clinical annotations were obtained from Maynard et al (*53*) which was analyzed using the Python package scanpy (*85*). The Kaplan-Meier curves displaying the overall survival of LUAD and PAAD patients according to the gene expression of the AXL downstream signature were generated using the web server GEPIA2 (*86*) which uses TCGA/GTEx data.

### Bacterial Display Library Preparation

The sequences used to construct the ‘Mass Spectrometry’ library were derived from peptide sequences of key proteins detected by mass spectrometry. Suspected phosphorylation sites were identified from these proteins to create 11-residue peptide sequences, with 5 residues flanking a central tyrosine on each end. The central tyrosine was represented with a ‘TAT’ codon, and the surrounding peptide sequences converted into DNA sequences that are codon-optimised based on E. coli codon usage and avoiding any SfiI restriction sites. For peptides containing tyrosine residues surrounding the phosphoacceptor tyrosine site, we substituted those surrounding tyrosine residues for phenylalanine. All sequences were then flanked with 5’-GCTGGCCAGTCTGGCCAG-3’ on the 5’ side and 5’-GGAGGGCAGTCTGGGCAGTCTG-3’ on the 3’ side, as oligos. This oligo pool was generated using on-chip massively parallel synthesis (Twist Bioscience), and PCR amplified with Oligo-pool forward (5’-GCTGGCCAGTCTGGC-3’) and reverse (5’-CAGACTGCCCAGACTGC-3’) primers over 10 cycles. All PCR reactions were done with Q5 high-fidelity DNA polymerase (NEB), and PCR products were subsequently purified with DNA clean-up after each PCR and restriction. DNA clean-up and gel extractions were performed with QiAquick Gel Extraction Kit (Qiagen).

This library is constructed with the pBAD33 plasmid vector, with an eCPX displaying gene tagged with c-Myc tag. The eCPX scaffold was PCR amplified with eCPX forward (5’-GGAGGGCAGTCTGGGCAGTCTG-3’) and eCPX reverse (5’-GGCTGAAAATCTTCTCTCATCCGCC-3’) primers to create a scaffold. This PCR product underwent a second PCR with the same eCPX Reverse primer, but with the oligo-pool library as the forward primer. The final PCR product library contained two SfiI restriction sites.

Simultaneously, the pBAD33 plasmid containing two SfiI restriction sites flanking HindII and PstI restriction sites underwent SfiI digest at 50℃ for 1 hour to generate the backbone. The reaction was subject to additional HindII and PstI digest at 37℃ for 1 hour to remove undesired insert by-products and undigested vector from the SfiI digest. The backbone was obtained by gel purification. The PCR product was subject to SfiI digest at 50℃ for 1 hour, followed by DNA clean-up to generate the insert.

The purified library insert was ligated into the digested pBAD33-eCPX backbone using T4 DNA ligase (NEB) for 1 hour at 25℃. The ligation involved 50 ng of DNA, with a 1:3 molar ratio of backbone:insert. The ligation reaction was used to transform commercial MC1061F cells by electroporation and purified using the QIAprep Spin Miniprep Kit (Qiagen).

### Preparation of E. Coli for Peptide Display of different Peptide Libraries

25 µL of electrocompetent E. coli SS320 (MC1061 F’) cells were transformed with 200 ng of library DNA, and the cells recovered with 1 mL of warm LB at 37℃ for 1 hr with shaking. These cells were resuspended in 250 mL of LB / chloramphenicol and incubated overnight at 37℃. 90 μL of the overnight culture was used to inoculate 5 mL of LB / chloramphenicol. This culture was grown at 37℃ for 1–2 hr until the cells reached an optical density of 0.5 at 600 nm, where 0.2% Arabinose was introduced for the induction of eCPX and peptide libraries. The culture was incubated for 4 h at 25℃. Afterwards, small aliquots (75 μL) of the culture were pelleted via centrifuge at 4000 g for 15 min, and the pellets were resuspended with 50 μL of kinase screen buffer (50 mM Tris, 10 mM MgCl_2_, 150 mM NaCl, 2 mM Na_3_VO_4_ and 1 mM tris (2-carboxyethyl) phosphine).

### Evaluation of AXL Library Phosphorylation by Flow Cytometry

AXL kinase was obtained commercially from Carna Bioscience, as GST-AXL (GST-Tagged) and BTN-AXL (Biotin Tagged). 2.7 μM of the kinase was subject to auto-phosphorylation with 5 mM of ATP and 10 mM MgCl_2_, for one hour at room temperature. 5 μL of the auto-phosphorylated kinase was added to 50 μL of resuspended cells displaying peptides on eCPX. The mixture was heated to 37℃ for 2 minutes, and 0.5 μL of 100 mM ATP was added to the reaction mixture. To evaluate AXL kinase activity 10 μL aliquots were removed from the reaction mixture at 5-, 10-, 20-, 40-, and 80-minute intervals. For the final assay, 30 minutes reaction time was used. The reaction mixtures were quenched with introduction of 25 mM EDTA to aliquots and cooling at 4℃. Subsequently, the cells were pelleted at (1500 g, 20 min), and washed with 20 μL PBS with 0.2% bovine serum albumin (BSA), before being pelleted as a precursor to antibody incubation. To evaluate the efficiency and kinetics of the phosphorylation reaction, the 10 μL pellets from 5-, 10-, 20-, 40-, and 80-minute reaction intervals were labelled by resuspension with 1:25 dilution of the PY20-PerCP-eFluor 710 conjugate (eBioscience) and 9E10 Alexafluor-488 conjugate (eBioscience) in PBS containing 0.2% BSA for 1 hour on ice, in the dark. These antibodies revealed the level of phosphorylation and myc (control, reference epitope) expression respectively. The cells were subsequently pelleted, and washed again with PBS with 0.2% BSA, before being diluted in 500 μL of PBS with 0.2% BSA for subsequent analysis using flow cytometry (BD LSR-Fortessa Cell Analyzer). To achieve similar library kinases phosphorylation levels as described previously (*61*), an optimal concentration of kinase and reaction time was determined to achieve around 30% library phosphorylation (**Fig. S4G-I**).

### Magnetic Beads Pulldown

100 μL of cells phosphorylated by AXL kinase (both GST-AXL, BTN-AXL) for 30 minutes was quenched and washed as described in ‘Preliminary assays to evaluate AXL phosphorylation’. The pelleted cells were resuspended on ice with 100 μL of 0.2% BSA, 1 mg/mL (1:500 dilution) of biotinylated anti-phosphotyrosine 4G10 (Merck) antibody and incubated for 1 hour. The cells were washed and resuspended with 0.1% BSA, 2 mM EDTA in PBS (Isolation buffer).

Following this, streptavidin beads (Dynabeads FlowComp Flexi Fit, Thermo Fisher) were rinsed with 1 mL of isolation buffer. 450 μL of isolation buffer were added to 50 μL of the antibody-labelled cells, and 75 μL of washed streptavidin beads were added. The mixture was rotated for 30 min at 4℃. Another 375 μL of isolation buffer was added to the beads-cell mixture before the beads were pulled down via a magnetic rack. The supernatant was removed, and the remaining beads were resuspended in 1 mL of isolation buffer and rotated for 30 min at 4℃. The beads were pulled down again with a magnetic rack and resuspended in 100 μL of water. These beads were boiled for 10 mins at 100℃ to lyse the cells. The mixture was centrifuged to pellet the beads, and the supernatant was extracted to obtain DNA for deep sequencing. In addition, 50 μL of cells from the PBS resuspension which were not labelled with antibodies were also boiled for 10 mins at 100℃ to obtain DNA that was input into the cells.

### Deep Sequencing and enrichment detection

20 μL of the lysate DNA containing the peptide library was PCR amplified with Q5 high-fidelity DNA polymerase (New England Biolabs), for 15 cycles in a 50 μL reaction, with Lib Prep 1 Forward (5’-TGACTGGAGTTCAGACGTGTGCTCTTCCGATCTNNNNNNNGCAGGTACTTCCGTAGC-3’) and Lib Prep 1 Reverse (5’-CACTCTTTCCCTACACGACGCTCTTCCGATCTNNNNNNNTTTTGTTGTAGTCACCAGA CTG-3’). 0.33 μL of the PCR product was then used for a second PCR reaction to attach the TruSeq UDI (Illumina) indexes, via a 20 cycle 50 μL PCR reaction with Q5 high-fidelity DNA polymerase (NEB), using Lib Prep 2 Forward (5’-CAAGCAGAAGACGGCATACGAGATGTGACTGGAGTTCAGACGTG-3’) and Lib Prep 2 Reverse (5’-AATGATACGGCGACCACCGAGATCTACACACACTCTTTCCCTACACGAC-3’) oligos. The resulting PCR product was purified using SPRIselect beads (Beckman Coulter). The samples were then submitted to the for Miseq and Novaseq deep sequencing (Illumina). Two technical replicates were performed on cells from the same library transformation, and two biological replicates were performed on cells from different library transformations, done on different days.

### Analysis and software

FLASH software was used to merge pair-end reads, and cutadapt was used to remove flanking sequences surrounding the variants of interest. To calculate enrichment scores of each peptide variant, the following equation was used.

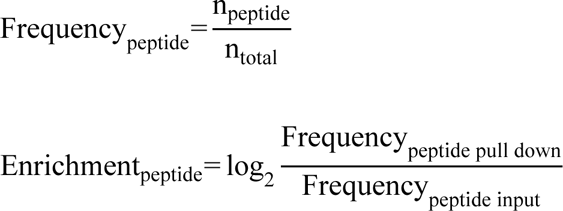

The number of reads of each peptide of each sample (*n*_*peptide*_) was first normalised as a frequency value *Frequency*_*peptide*_to the total number of reads for every peptides encoded by the library (*n_total_*. Then, *Frequency*_*peptide*_ was compared between the pull down (*Frequency*_*peptide pull down*_) and input (*Frequency*_*peptide input*_), as a log 2-fold change ratio. To compare enrichment scores between replicates, the z-score of each peptide’s enrichment was calculated, using:

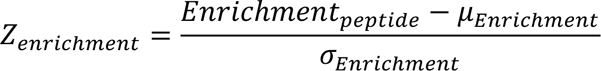

Where σ_*Enrichment*_ is the standard deviation of the library’s enrichment score of the replicate, and μ_*Enrichment*_ is the standard deviation of the library’s enrichment score.

### Drug-Tolerant Persister Cell Assay

1,000 cells were seeded in a 96-well plate and then treated with the indicated treatments and cell lines, with four technical replicates per treatment. After 15 days, the treatment solutions were replaced with complete RPMI-1640 media for an additional 15 days to allow drug-tolerant persister cells to regrow. Cells were imaged every 4 hours using an IncuCyte S3 (Essen Bioscience) at 10x magnification with 4 images per well. Phase images were manually thresholded and then analyzed by IncuCyte S3 software (Essen Bioscience) to determine cell confluency.

## Acknowledgements

We thank A. Li and N. Shah for sharing reagents, the pTyr-Var library and their advice setting up peptide display screens, as well as the Cancer Research UK Cambridge Institute core facilities for the deep sequencing and flow cytometry analysis of our screens.

## Author contributions

A.S.M conceived the study. A.S.M. and M.C led the study. F.W.M and P.C contributed to the refinement of the study. S.Y.B and S.D.T constructed the PC9 AXL mutant cell lines. M.C and S.D.T performed the phenotypic measurements of the PC9 AXL Y-to-F mutants, performed sample preparation for mass spectrometry experiments, biochemical assays including western blots, and ELISA-based Luminex, as well as drug-tolerant persister assays. M.L and C.B contributed to the experiments. M.C performed computational analyses, mathematical modeling, and immunofluorescence experiments. S.D.T generated samples for RNA-seq. J.G performed all the mass spectrometry-based phosphoproteomics experiments. P.C, M.J, and T.A devised and performed the bacterial display specificity screen of AXL. A.S.M and F.W.M secured funding.

## Data and materials availability

All analysis was implemented in Python and can be found at https://github.com/meyer-lab/AXLomics.

