## Supplementary Figures for "Dissecting signaling regulators driving AXL-mediated bypass resistance and associated phenotypes by phosphosite perturbations"

### SUPPLEMENTARY MATERIALS

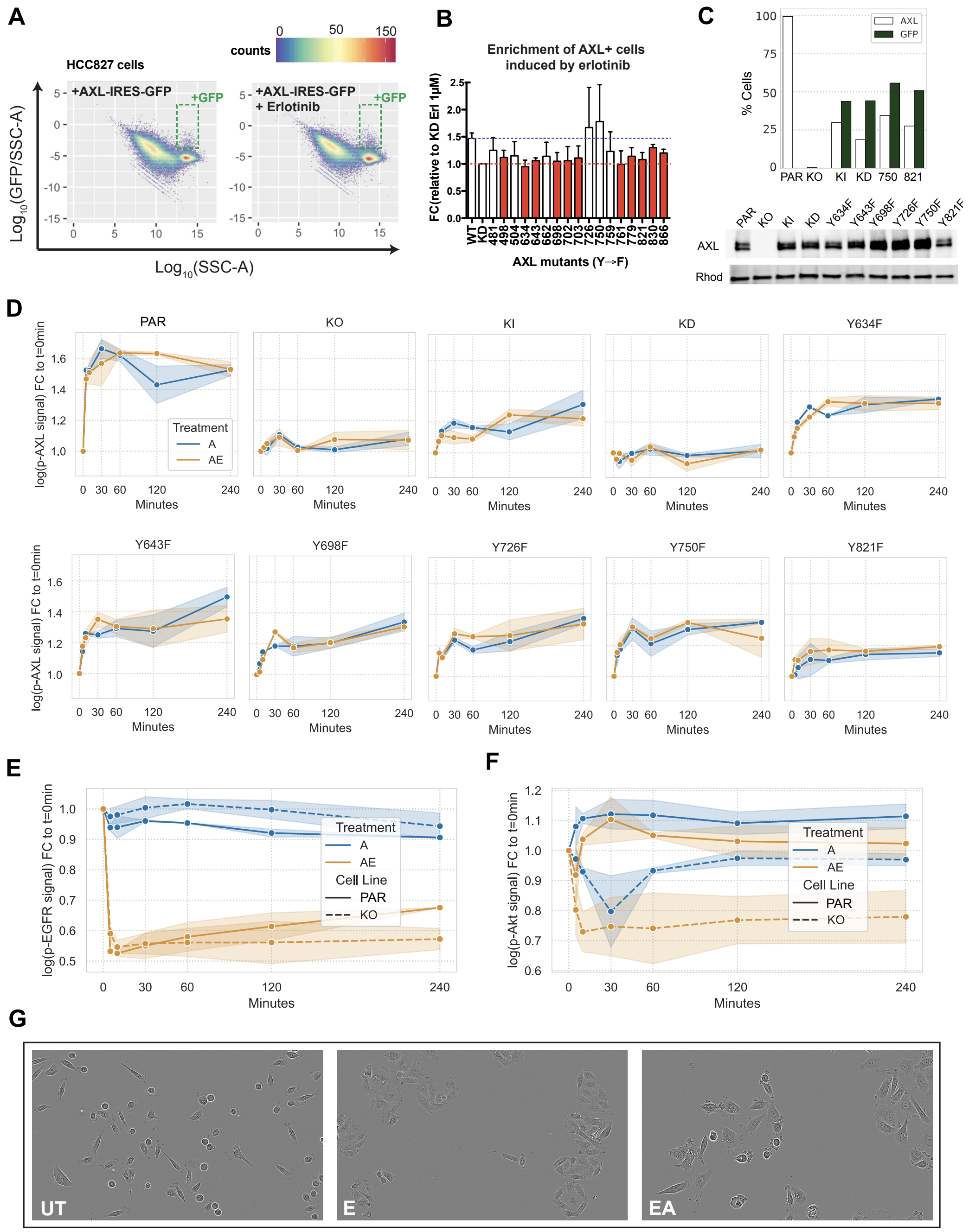


**Figure S1. AXL expression and activation across PC9 AXL Y-to-F mutant cell lines.** (A–B) AXL KO HCC827 cells were transfected with an AXL (Y-to-F)-IRES-GFP cassette (A) to quantify the enrichment of AXL+ cells induced by E (B). (C) Cell surface and total protein AXL expression in the indicated PC9 cell lines measured by either FACS (top) or western blotting (bottom). (D) AXL phosphorylation signal in PC9 AXL mutant cell lines in response to A alone or in combination with E. Error bars show the standard error of the mean. (E) EGFR inhibition in response to E. (F) AXL-specific activation of Akt. (G) Images of the E-induced island effect in PC9 parental cells. In D–F the error regions correspond to the standard error of the mean. Related to **Figure 1**.

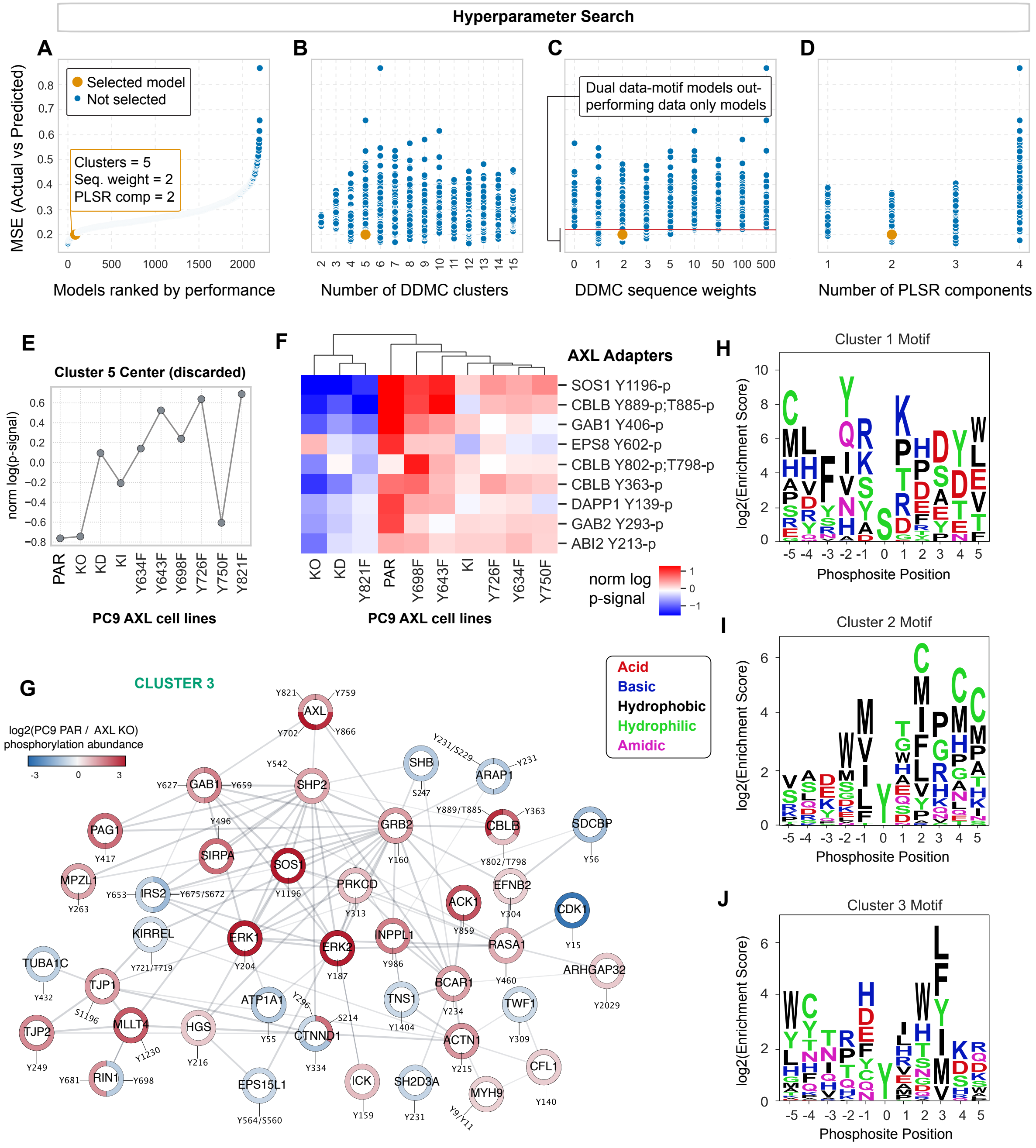


**Figure S2.** (A–D) Hyperparameter search based on the ability of DDMC centers to predict AXL-mediated phenotypes using PLSR. Mean-Squared Error (MSE) of actual versus predicted phenotypic measurements grouped by model performance (A), number of DDMC clusters (B), DDMC sequence weights (C), and number of PLSR components (D). Note that even though some models were slightly more predictive, we selected the model using 5 clusters, a sequence weight of 2, and a 2 PLSR components due to its interpretability. (E) DDMC Cluster 5 center which was discarded due to biologically inexplicable signaling behavior. (F) Hierarchical clustering of the phosphorylation signal of RTK adapters across AXL Y-to-F mutants. (G) STRING network of DDMC's C3. (H–J) Logo plots illustrating the PSSMs of each cluster. Related to **Figure 2**.

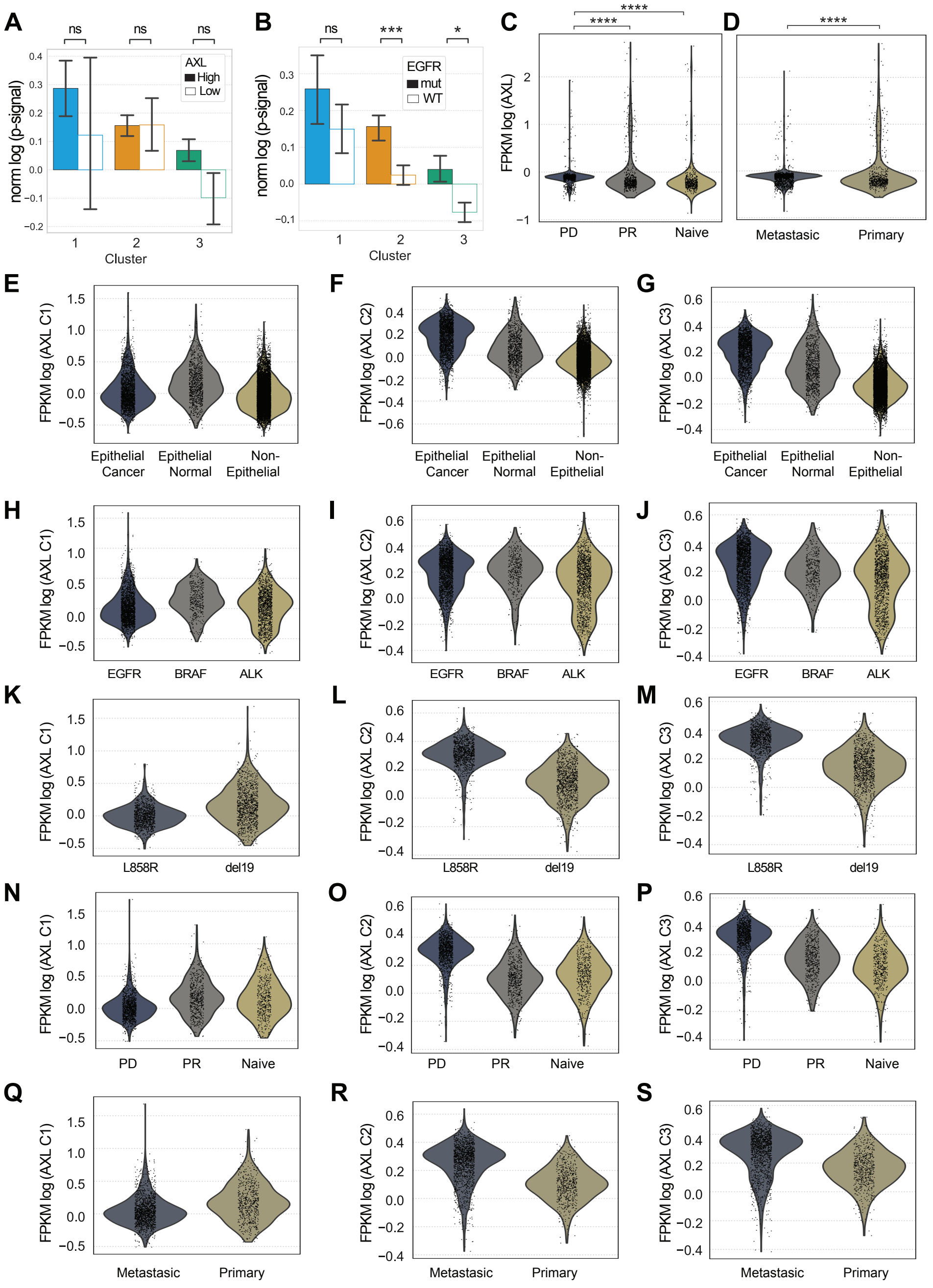


**Figure S3** (A–B) Phosphorylation signal of C1, C2, or C3 in AXL-hi and AXL-low (A) and EGFRm or EGFR WT tumors (B). Error bars in represent the standard error of the mean. (C–D) AXL expression grouped by treatment response (C, PD: progressive disease, PR: partial response) or metastatic status (D). (E–S) Gene expression of C1, C2, C3 in (E–G) epithelial cancer cells, epithelial normal cells, or non-epithelial; based on (H–J) gene driver, (K–M) EGFR mutation, (N–P) tumor response, and (Q–S) metastasis status. Statistical significance in A-D was calculated by a Mann-Whitney U rank test. *p-value < 0.05, ***p-value < 0.0001, and ns means not significant. Related to **Figure 3**.

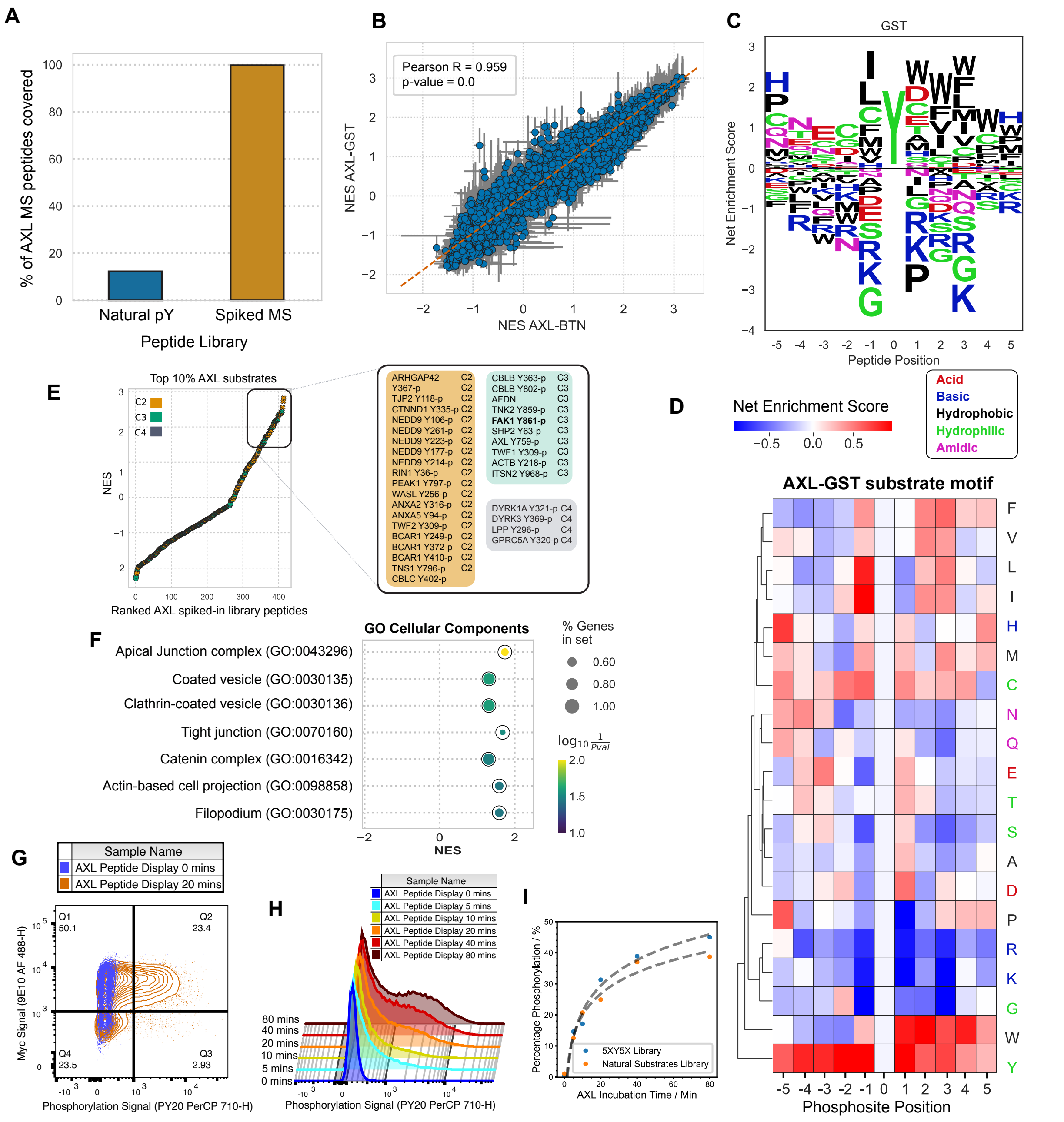


**Figure S4.** (A) Percentage of peptides covered by either the pTyr-Var library or after spiking with AXL-regulated peptides identified in phosphoproteomic analysis. (B) Pearson correlation of NES between AXL-GST and AXL-BTN. (C–D) AXL PSSM motif illustrated by either a logo plot (C) or a heatmap (D). (E) Ranked AXL substrates by NES and colored by cluster membership, alongside a selected list of phosphosites per cluster. (F) Ranked GSEA using the

GO Cellular Components gene set. (G) Scatter plot of myc and phosphorylation signal of peptide display library in samples incubated during 0 or 20 minutes. (H) Histogram showing phosphorylation levels of peptide display library during incubations of 0, 5, 10, 20, 40, or 80 minutes. (I) Phosphorylation percentage as a function of time using two different peptide libraries (see methods for details). Related to **Figure 5**.

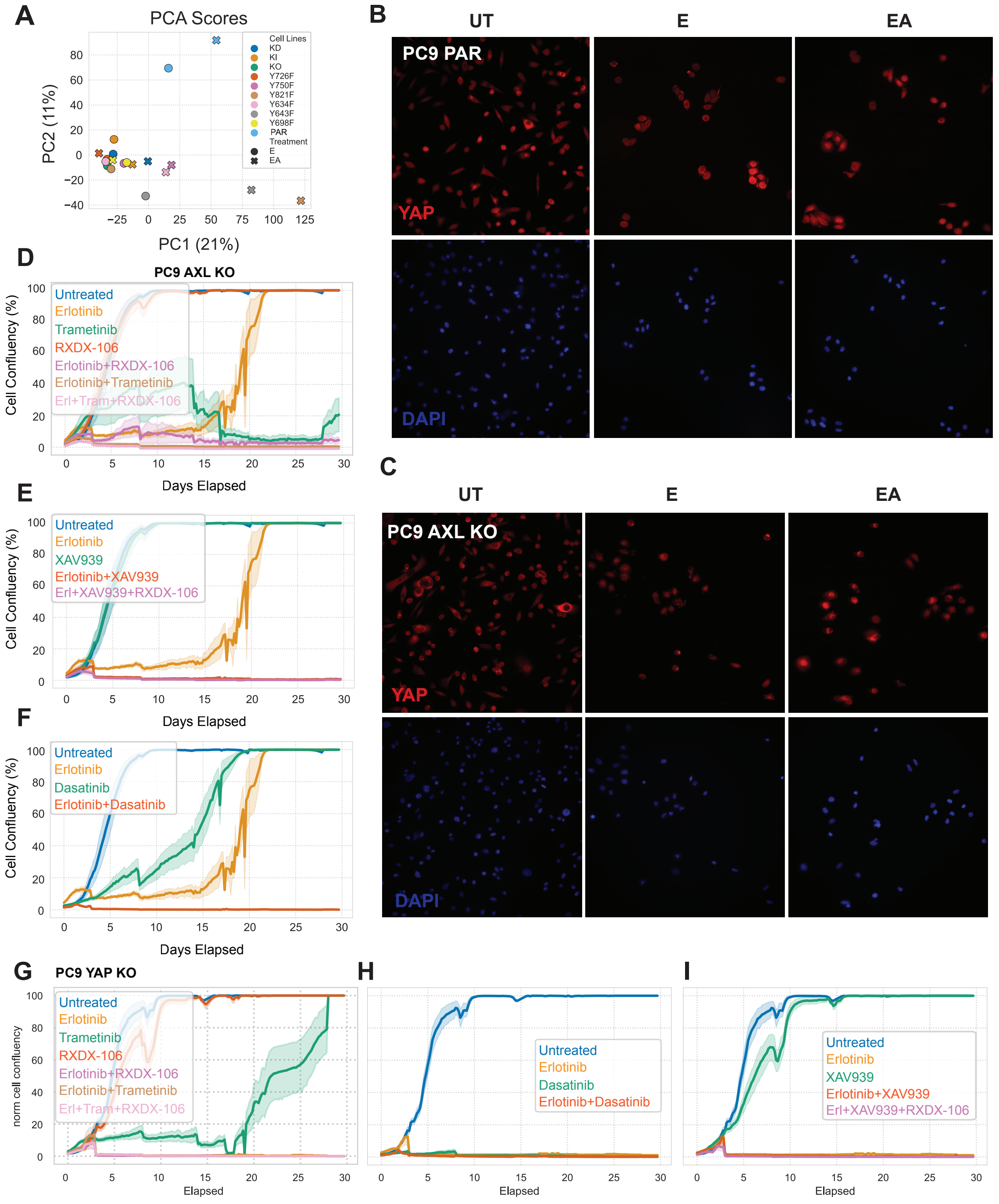


**Figure S5.** (A) Scores of a PCA analysis of the RNAseq data of the PC9 Y-to-F mutant cell lines treated with E or EA. (B–C) Immunofluorescence staining of YAP and DAPI in (B) PC9 parental and (C) AXL KO cells. (D–I) Cell viability assay of PC9 AXL KO cells (D–F) or YAP KO cells (G–I) treated with the indicated inhibitors for 15 days. Treatment solutions were then replaced with media and drug-tolerant persister cells were allowed to regrow for 15 days. Error regions show the standard error of the mean.
